# Signatures of covert neuron loss in the local field potential of motor cortex

**DOI:** 10.64898/2025.12.09.693286

**Authors:** Kenji Marshall, Stephen E. Clarke, Paul Nuyujukian

## Abstract

**Background:** Covert stroke is understudied despite occurring ten times for every symptomatic stroke and contributing to stroke’s enormous global disease burden. For instance, it is unknown whether covert stroke replicates the peri-infarct neuroelectrophysiological frequency spectrum impact of symptomatic stroke. This motivated the use of our novel electrolytic lesioning platform to explore the spectral consequences of covert neuron loss.

**Methods:** During a multi-month period of participation in an arm reaching task, neuron loss was induced via electrolytic lesions to the motor cortex of two large animals (U: *n* = 4; H: *n* = 7). Behavioral metrics were paired with spectrum estimates from local field potential recordings. These spectra were represented as bandpowers, aperiodic/periodic parameters, and decomposed time-frequency tensors. Lesion impact was assessed using nonparametric permutation tests of next-day effects and state space model parameters.

**Results:** Task success rate was unaffected by lesions, but shifts in aperiodic structure reduced next-day *γ* bandpower (30 – 100 Hz; U: −1.38µV^2^, *p <* 1 × 10*^−^*^3^; H: −1.66µV^2^, *p* = 0.001) while sensorimotor rhythms spanning 8 – 45 Hz (Σ*SMR*) were amplified (Monkey U: 8.68µV^2^, *p <* 1 × 10*^−^*^3^; Monkey H: 2.40µV^2^, *p* = 0.004). Additionally, state space modeling showed that spectral perturbations to *γ* and Σ*SMR* outlasted any behavioral impact of the lesions (Monkey U: Behavior=1 d, *γ* =2 d, Σ*SMR* =2 d; Monkey H: Behavior=0 d, *γ* =3 d, Σ*SMR* =1 d). Finally, tensor decomposition revealed interpretable, animal-specific perturbations to time-frequency dynamics.

**Conclusion:** Covert neuron loss induced multi-day spectral perturbations similar to those observed after symptomatic stroke. The neural spectrum is thus more sensitive to neuron loss than previously understood and could be responsive to neuron loss caused by covert stroke. This work also motivates the use of electrolytic lesions to bridge covert and symptomatic regimes of neuron loss, advancing our causal understanding of post-stroke interactions between spectrum and behavior.

## Introduction

Electrophysiological recordings of the peri-infarct cortex have shown that ischemic stroke induces measurable changes in the neural spectrum, correlated with behavioral recovery and stroke severity.^1–23^ These spectrum perturbations may provide a window into how functional networks reorganize after stroke^24–31^ and could serve as quantitative rehabilitation targets.^32^ But, accessing this scientific and translational impact requires clarity on the causal relationships between stroke, spectrum perturbations, and behavioral outcomes. For instance, if we assume that the causal structure can be represented as a graph–with the underlying stroke as the root–we cannot easily distinguish between the graphs depicted in Figure 1A. Is the spectrum perturbation caused by behavioral deficit or vice versa? Are the two effects generic, independent responses to stroke of a given scale? Of course, these graphs simplify a reality which is much more complex and dynamic, but the point remains: a clinically relevant theory of the post-stroke neural spectrum requires mapping out this causal terrain.

**Figure 1:**
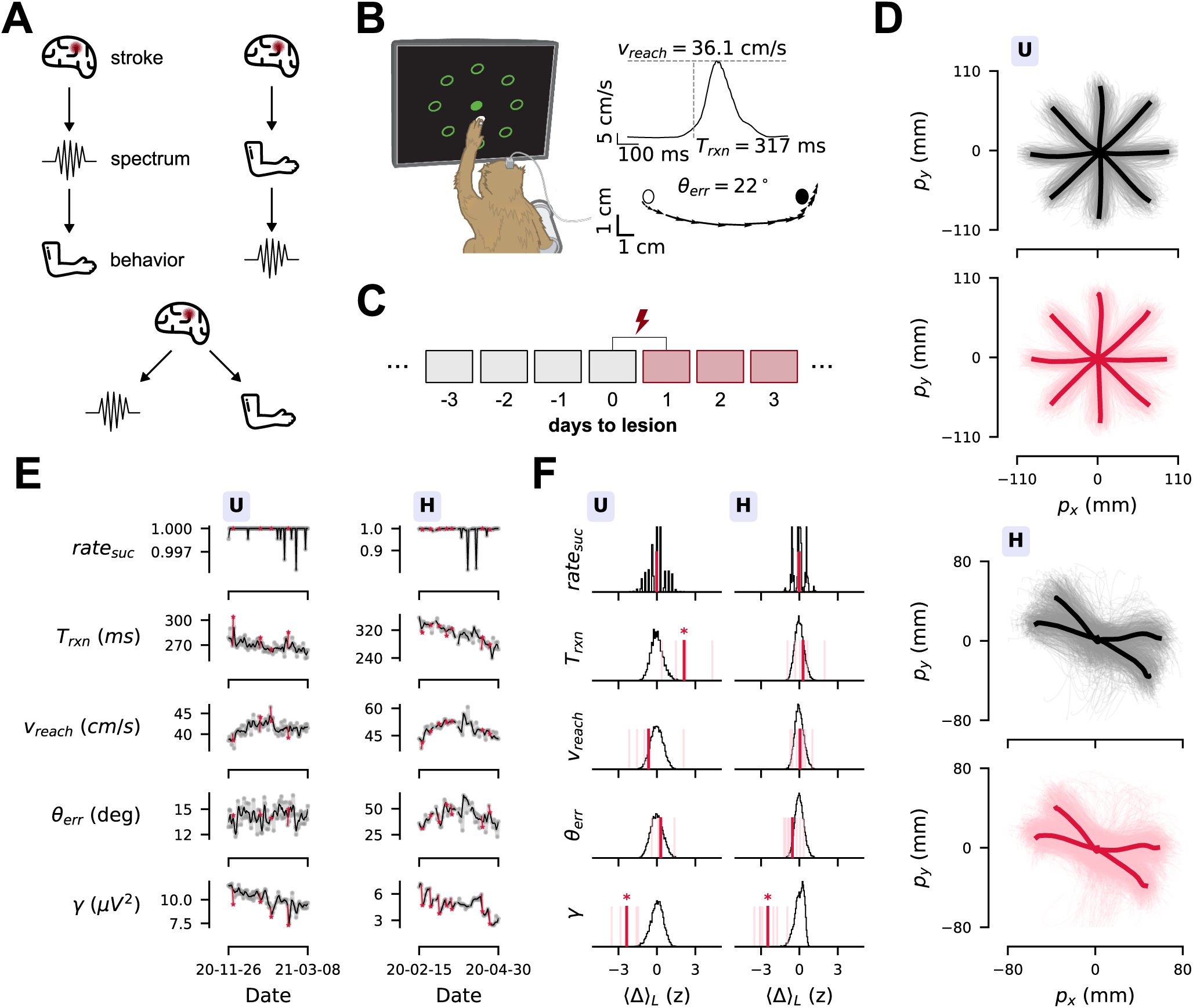
Lesions were covert but perturbed bandpower features. **(A)** Causal possibilities for post-stroke effects on behavior and the neural spectrum are difficult to distinguish from correlational studies reporting behavioral breakdown alongside spectrum perturbation. **(B)** Monkeys U/H participated in a center-out reaching task, where individual trials were kinematically represented by a reaction time (*T*_rxn_), reach speed (*v*_reach_), and angular error (*θ*_err_). **(C)** An electrolytic lesioning platform^53^ induced covert neuron loss in the midst of longitudinal recordings (Monkey U: *n* = 4, Monkey H: *n* = 7) via the implanted microelectrode array. Lesions occurred after recording on Day 0, then neural activity and behavior were sampled ∼24 hours later on Day 1; this was repeated multiple times. **(D)** Kinematics were qualitatively preserved after lesions, as shown by visualizing cursor position across Day 0 (black) and Day 1 (crimson) reaches. Thin traces indicate single reaches, while thick lines indicate mean trajectory for each reaching target. Monkey U played a traditional 8-target task while Monkey H played with 6 targets, 4 of which produced stereotyped reaches (see Detailed Methods, Figure S1A). **(E-F)** Lesions were embedded in longitudinal recordings (E), while permutation tests of a lesion sensitivity statistic ⟨*Δ*⟩*_L_* (Detailed Methods) determined significant next-day impacts from lesions (\**p <* 0.05) (F). In (E), the Day 0 to Day 1 effects are marked in crimson, while the black trace is a smoothed version (Detailed Methods) of the raw gray data. Notably, task success rate (rate_suc_) was unaffected (Monkey U: ⟨*Δ*⟩*_L_* = −0.01, *p* = 1.000; Monkey H: ⟨*Δ*⟩*_L_* = −0.02, *p* = 0.828), suggesting that neuron loss was covert. Other behavioral metrics were also unaffected except for *T*_rxn_ in Monkey U (⟨*Δ*⟩*_L_* = 2.12, *p <* 1 × 10*^−^*^3^). However, both monkeys experienced a significant reduction to *γ* power after lesions (Monkey U: ⟨*Δ*⟩*_L_* = −2.36, *p <* 1 × 10*^−^*^3^; Monkey H: ⟨*Δ*⟩*_L_* = −2.47, *p* = 0.001).See all lesion sensitivity statistics in Table S2).

To resolve (and improve upon) the structures in Figure 1A, measuring spectral impact at the boundary between covert stroke (defined as ischemia occurring “without a history of acute neurological dysfunction attributable to the lesion”^33^) and symptomatic stroke could unify spectral shifts related to behavioral compensation^34^ and those that are relevant to behavioral deficit. These measurements are also motivated by the prevalence and disease burden of covert stroke,^35^ which occurs relative to symptomatic strokes at a rate of ∼10:1^36,37^ (affecting up to 30 % of the elderly^38^) while increasing risk for dementia,^39,40^ depression,^41^ and future symptomatic strokes.^39,42–44^ Despite this, our scientific understanding of covert stroke is limited and there is “little evidence for how to investigate or manage” covert stroke in clinical contexts, as described in an official statement by the American Heart Association/American Stroke Association (AHA/ASA).^35^ There is thus a scientific and clinical need for basic animal models of covert stroke, the translational value of which would be maximized in the gyrencephalic primate brain due to anatomical similarities with humans in cortical organization and descending motor pathways.^45^ This need for primate stroke models^20,46–48^–that can connect the rodent stroke literature^25,26,29^ with human neurophysiology–has been reiterated for over 25 years by expert consortia^45,49–51^ and the US National Institutes of Health.^52^ Considering the unique ethical demands of working with non-human primates (NHPs), covert stroke is an appealing starting point.

In response to these needs, we used our novel electrolytic lesioning platform^53,54^ to model the functional consequences of covert stroke. Although this model cannot capture all aspects of covert stroke’s vascular origin and molecular/cellular pathology, it intentionally isolates covert neuron loss as the core functional consequence of covert stroke. This allowed us to measure, for the first time, the spectral impact of covert neuron loss in two rhesus macaques (Monkeys U and H) while recording reach-related local field potential (LFP) data. We found that covert neuron loss, which did not affect behavioral metrics like success rate and kinematic entropy, induced spectrum perturbations that were coherent with the symptomatic stroke literature, including a prominent reduction to *γ* bandpower (30 – 100 Hz)^4,19–22^ driven by shifts in the aperiodic spectrum structure,^22,23,55–57^ alongside heterogeneous effects on movement-related time-frequency dynamics.^11–16^ The lesions also increased the power of sensorimotor rhythms (SMRs) in the 8 – 45 Hz frequency range. Overall, we motivate the use of controlled neuron loss in NHPs across multiple scales of injury in order to functionalize the neural spectrum as a tool for understanding and treating stroke, across covert and symptomatic regimes.

## Methods

Additional details on the following methods are available in the Supplemental Material.

### Data availability

Derivative datasets and associated Python code to reproduce core results will be made available on the Stanford Digital Repository (deposition pending).

### Animal care

Two adult male rhesus macaques (*Macaca mulatta*), aged 11 (Monkey U) and 14 (Monkey H), were implanted with 96-channel Utah arrays in the arm region of contralateral primary motor cortex (M1) (Blackrock Microsystems, Salt Lake City, UT, USA). All animal procedures were reviewed and approved by Stanford University’s Institutional Animal Care and Use Committee.

### Behavior

Subjects performed a center-out reaching task for a juice reward (Figure 1B). Alongside a daily success rate (rate_suc_), reach kinematics were optically tracked and the daily medians of reaction time (*T*_rxn_), reach speed (*v*_reach_) and angular error (*θ*_err_) were computed (Figure 1B). Trial inclusion criteria ensured reaches were stereotyped (Detailed Methods, daily rate of trial inclusion denoted rate_acc_), and then electrophysiological data was extracted from epochs spanning 150 ms before to 350 ms after *T*_rxn_.

To supplement the standard behavioral features of rate_suc_, *T*_rxn_, *v*_reach_, and *θ*_err_, a kinematic entropy measure was used where histograms of daily 2D arm position and velocity were constructed and then entropy was computed to yield *H_p_* and *H_v_*.

### Electrolytic lesions

The lesion circuit^53^ used pairs of adjacent electrodes as the anode/cathode to pass controlled amounts DC current through tissue. At an amplitude of 150 – 170 µA to M1, Monkey U received four 30 s lesions, while Monkey H received ten lesions mostly at a duration of 45 s (lesion three was 130 µA for 50 s while lesion ten was 480 µA for 30 s) (Table S1). This parameter range was calibrated using histology in *ex vivo* ovine and porcine brains and *in vivo* anesthetized porcine and macaque brains^53^ to evoke lesion volumes on the order of ∼1 mm^3^, a fraction of ∼20 mm^3^ photothrombotic^20^ or ∼100 mm^3^ aspiration^58^ lesions previously reported in macaque sensorimotor cortex. Due to recording issues in lesion 1 and non-participation after lesion 3, only the last 7 lesions of Monkey H featured consistent daily recordings and were included for analysis. Blood testing, ear temperature, neurological/physical examinations, and extensive in-cage monitoring were used to ensure animal well-being after lesions; no abnormalities were documented (Table S1).

### In vivo electrophysiology

Daily periodograms were estimated from filtered and downsampled 1 kHz LFP data between 1 – 100 Hz by taking a median across all channels and trials for a given day; canonical bandpowers were then computed (*δ*: 1 – 4 Hz,*θ*: 4 – 8 Hz, *α*: 8 – 12 Hz, *β*: 12 – 30 Hz, *γ*: 30 – 100 Hz). A time-frequency tensor (Figure 4A) was formed using squared wavelet coefficients (complex Morlet;^59,60^ center frequency: 1 Hz; bandwidth: 2 Hz^61^) from 8 – 100 Hz in increments of 1 Hz and in 10 ms temporal increments from −320 ms to 120 ms relative to *T*_rxn_.

For daily periodogram data, spectral parametrization^55^ hyperparameters were selected (Figure S3B), yielding an aperiodic exponent (*ap*_exp_) and offset (*ap*_off_), alongside the locations, heights, and widths of oscillatory peaks. Two recurring SMRs in the *α*/*β* range were identified where possible (*SMR_↓_*, *SMR_↑_*) alongside their powers (*SMR_↓_^P^*, *SMR_↑_^P^*), frequencies (*SMR_↓_^f^*, *SMR_↑_^f^*), and widths (*SMR_↓_^W^*, *SMR_↑_^W^*). An aggregate SMR power was estimated each day from 8 – 45 Hz, denoted Σ*SMR*.

In addition, the temporal dynamics of average wavelet coefficients in 8 Hz bands around median *SMR_↑_^f^* were analyzed (Monkey U: 29 – 37 Hz; Monkey H: 31 – 39 Hz), and a pre-movement baseline (*β*_ref_) was computed alongside a normalized movement-related beta decrease (MRBD) and post-movement beta rebound (PMBR). Next, the nonnegative canonical polyadic decomposition (CPD) reduced the time-frequency tensor to a sum of rank-one component tensors.^62–65^ A rank *R* CPD provided *R* factors across days (*τ_r_^d^*), frequency (*τ_r_ ^f^*), and time (*τ_r_ ^t^*) which approximated the original tensor (Figure 4A). Ranks of *R* = 4/*R* = 3 were selected for Monkeys U/H.

### Statistics

To identify next-day lesion effects, *z* -scored first differences of aforementioned features between the pre-lesion recording session (Day 0 in Figure 1C) to another session ≈24 h later (Day 1 in Figure 1) were averaged, yielding the standardized statistic ⟨*Δ*⟩*_L_*. Next, a linear Gaussian state space model (LG-SSM) was used as an alternate model of lesion impact that also measured its temporal extent.^66^ Given a *z* -scored time series sampled at times *t* (*y_t_*), we modeled this as having three components: a component arising from a linear dynamical system (*x_t_*), a component that was driven by inputs (*u_t_*), and a noise component (Figure 3A). The lesions were modeled as the input *u_t_*, capable of influencing the time series for 5 days after a lesion occurred. To do so, *u_t_* was a 5-vector where the *i* th index was 1 when it was *i* days after a lesion and 0 otherwise (Figure 3A). The strength of this influence was captured by the standardized lesion input coefficients *d*_1_, *d*_2_, *. . .*, *d*_5_. Three such models were fit to *z* -scored Behavior, Periodogram, and Tensor datasets; for each feature, the marginal log-likelihood (SSM_LL_) and lesion input coefficients (*d*_1_, *d*_2_, *. . .*, *d*_5_) were analyzed for significance.

Significance of ⟨*Δ*⟩*_L_*/SSM_LL_/*d_i_* statistics was determined using nonparametric within-subject permutation tests,^67,68^ using the null hypothesis that observed data would be equally likely if lesion days had been randomly re-assigned. This approach means that monkeys acted as their own statistical control, where lesion impact was measured against empirical fluctuations in recorded longitudinal data. This permitted adherence to the systems neuroscience ethical standard of 2 monkeys. The false discovery rate was used to control the family-wise error rate at 0.05 for each group of tests.^69^ Beyond considering statistical significance, the ⟨*Δ*⟩*_L_*/*d_i_* statistics report a standardized impact of lesions on the relevant feature, providing an effect size interpretable in feature standard deviations. These statistics are also disclosed in natural units (e.g. µV^2^) in the relevant Supplemental Tables. See the Detailed Methods for additional statistical descriptions, including approaches for behavioral correlation, spatial correlation, and recurrent lesion effects.

## Results

### Covert lesions perturbed bandpower features

Monkeys U and H performed a center-out reaching task almost every day for months (Monkey U: 102/103 days; Monkey H: 70/76 days) during simultaneous LFP recordings from M1 (Figure 1B, left). Monkey U played a standard radial reaching task to eight targets, while Monkey H had the bottom and bottom left targets removed due to a visual deficit; in addition, reaches to the top and top right were poorly stereotyped and were discarded, leaving four reach conditions (Figure S1A, Detailed Methods). Up to 750 successful reaches were included in the dataset on each recording day yielding a total of 74,054 trials for Monkey U and 36,827 trials for Monkey H. In this period of time, electrolytic lesions were performed in M1 via the implanted array (Monkey U: 4 lesions, Monkey H: 7 lesions)^53^ (Figure 1C). Lesion parameters were selected to be covert, inducing neuron loss while preserving the ability of the animal to complete the arm-reaching task. By “covert”, we mean that the animal’s success rate (rate_suc_) was not impacted by neuron loss, in accordance with animal stroke models analyzing success rates in pellet reaching tasks^23,56,70–74^.

Near-identical reach kinematics were observed for trials before and after lesions (0 to 1 days to lesion in Figure 1C; kinematics shown in Figure 1D), suggesting a covert effect. To make this concrete, we leveraged the fact that lesion-induced fluctuations were sparsely embedded within longitudinal time series of neural recordings and behavior (crimson markings in Figure 1E). Thus, lesion perturbations for a given feature were compared to the empirical distribution of day-to-day fluctuations by permutation tests of the statistic ⟨*Δ*⟩*_L_* (mean, *z* -scored, lesion-induced fluctuation). Applied to rate_suc_, no lesion impact was detected (Monkey U: ⟨*Δ*⟩*_L_* = −0.01, *p* = 1.000; Monkey H: ⟨*Δ*⟩*_L_* = −0.02, *p* = 0.828; top of Figure 1F); in fact, rate_suc_ was over 99.5 % on all post-lesion days in each animal (top of Figure 1E). When considering other behavioral metrics, no impact was detected except for an increase to *T*_rxn_ for Monkey U (⟨*Δ*⟩*_L_* = 2.12, *p <* 1 × 10*^−^*^3^) (Figure 1E-F), amounting to a mean slowing of ∼17 ms. The rate at which trials were included for analysis based on passing behavioral filters (rate_acc_) was also unaffected (Figure S1B-C, complete ⟨*Δ*⟩*_L_* statistics in original and *z* -scored units are in Table S2).

To further validate the covert impact of the lesions, the behavioral metrics analyzed above were paired with the entropy of daily position/velocity kinematic distributions (Figure S2A). Reduction in movement entropy has a history of use in sports science as a summary statistics for tracking increased motor efficiency and skill throughout learning.^75,76^ Increased entropy of gait patterns has also been measured after acute peripheral ischemia-reperfusion.^77^ Thus, this measure offered a sensitive, data-driven supplement to our behavioral characterization. Position and velocity entropy (*H_p_* and *H_v_*) were computed each day (Figure S2B), and lesion impact was not observed for either *H_p_* (Monkey U: ⟨*Δ*⟩*_L_* = 0.03, *p* = 1.000; Monkey H: ⟨*Δ*⟩*_L_* = 0.64, *p* = 0.124) or *H_v_*(Monkey U: ⟨*Δ*⟩*_L_* = −0.26, *p* = 0.742; Monkey H: ⟨*Δ*⟩*_L_* = 0.15, *p* = 0.817) (Figure S2C, Table S2). In summary, across consideration of reaching ability (rate_suc_), standard metrics (*T*_rxn_, *v*_reach_, *θ*_err_) and entropy (*H_p_*, *H_v_*), the lesions had no impact besides an increase to Monkey U’s *T*_rxn_, in alignment with our definition of “covert”.

Contrasting the sparse behavioral impact, a robust *γ* power decrease on the order of 2 standard deviations was observed in both monkeys (Monkey U: ⟨*Δ*⟩*_L_* = −2.36, *p <* 1 × 10*^−^*^3^; Monkey H: ⟨*Δ*⟩*_L_* = −2.47, *p* = 0.001; bottom row of Figure 1E-F). Lesion impacts were also observed in other bands, such as an increase in *α* and *β* activity in Monkey U alongside a reduction in *δ*, *θ*, *α*, and *β* power in Monkey H (Figure S1B-C, Table S2). Interestingly, this pronounced loss of high-frequency power^4,19–22^ (or a relative increase in low-frequency power^2,3,5,7,8,78–81^) has been previously observed in symptomatic stroke. However, interpreting bandpower shifts is tenuous without differentiating between the periodic oscillations existing atop the aperiodic, power-law structure of the spectrum.^55,82^ The *γ* reduction observed here may be compatible with a shift in the aperiodic power law,^22,23,56,57^ while the heterogeneity in the *α* and *β* bands may relate to shifts in sensorimotor rhythms (SMRs) known to occupy those frequency ranges.^83–86^ As such, spectral parametrization was then applied.^55^

### Bandpower effects were driven by perturbed aperiodic/periodic parameters

After model selection (Figure S3B, Detailed Methods) and peak identification (Figure S3A), daily spectra were represented by their aperiodic parameters (*ap*_exp_, *ap*_off_) as well as the properties of oscillatory peaks; total periodic SMR power from 8 – 45 Hz was aggregated into the daily summary statistic Σ*SMR*. The parameters of two individual SMRs (*SMR_↓_*, *SMR_↑_*)–including power (*SMR_↓_^P^*, *SMR_↑_^P^*), frequency (*SMR_↓_^f^*, *SMR_↑_^f^*), and width (*SMR_↓_^W^*, *SMR_↑_^W^*)–were also analyzed (see Figure S3C-D). The model consistently had high goodness-of-fit (median *R*^2^ across days [IQR]; Monkey U: 0.9947 [0.9941, 0.9952]; Monkey H: 0.9977 [0.9960, 0.9987]) which was not affected by the lesions (Figure S3C-D). Monkey U showed a significant increase in *ap*_exp_ (⟨*Δ*⟩*_L_* = 1.61, *p* = 0.003) while *ap*_off_ was unaffected (Figure 2B-C). This perturbed aperiodic structure (Figure 2D, top) in a way compatible with *γ* power loss, as observed previously.^22,23,56,57^ In Monkey H, there was a marginally insignificant effect on *ap*_off_ (⟨*Δ*⟩*_L_* = −0.81, *p* = 0.079) (Figure 2B-C); resultant aperiodic shifts also visibly reduced *γ* power (Figure 2D, bottom). The close relationship between aperiodic activity and *γ* power is further supported by the fact that aperiodic *γ* power explained a large proportion of *γ* power variance (*R*^2^; Monkey U: 0.84; Monkey H: 0.89) while periodic *γ* power did not (*R*^2^; Monkey U: 0.02; Monkey H: 0.03).

**Figure 2:**
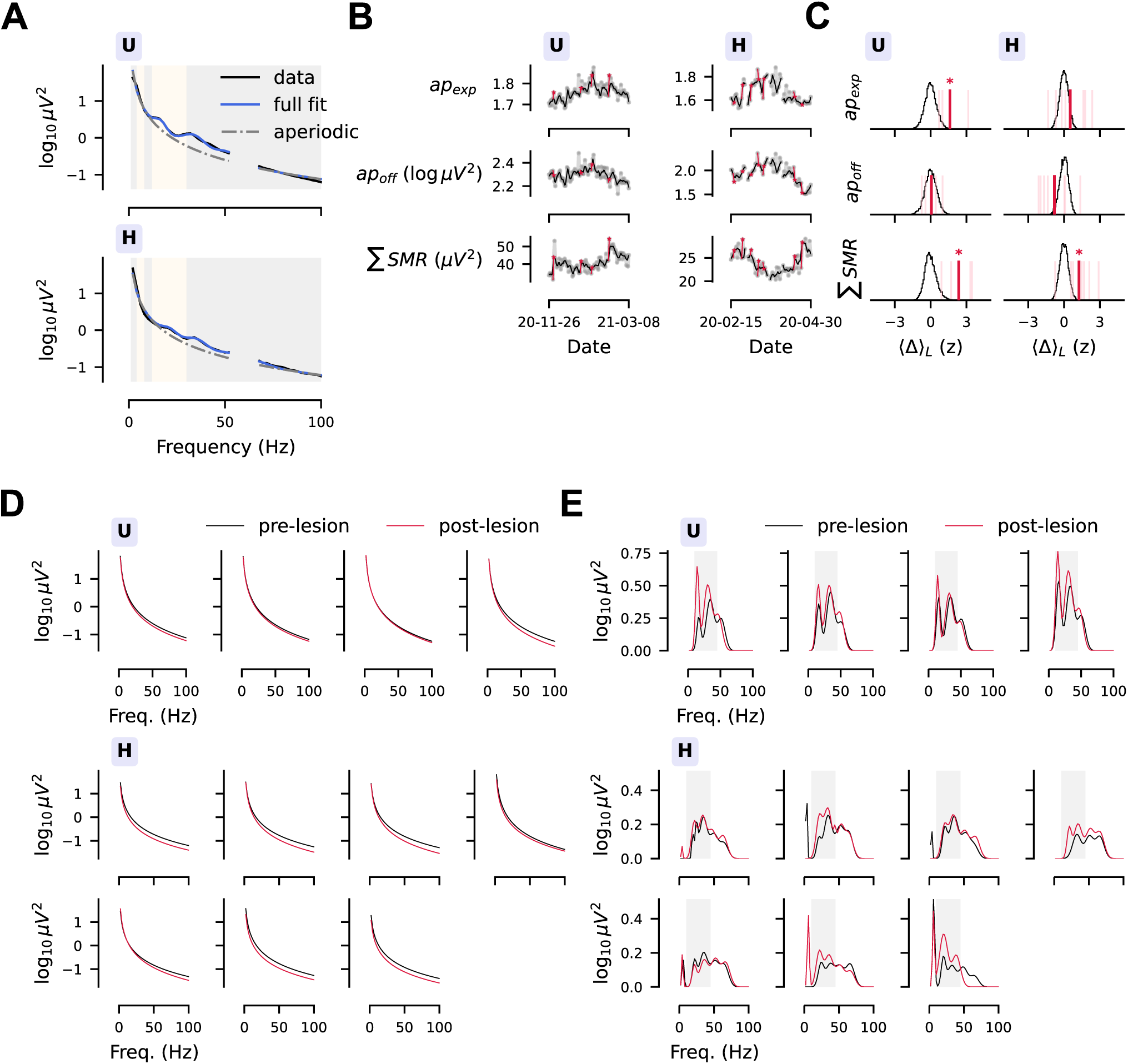
Bandpower effects were driven by perturbed aperiodic and periodic parameters. **(A)** Using spectral parametrization,^55^ daily periodograms (black) were represented as aperiodic (gray) and periodic (blue shows aperiodic + periodic) components. Shown are representative fits to Monkey U’s spectrum on 2020-12-01 (*ap*_off_ = 2.33 logµV^2^, *ap*_exp_ = 1.73, peaks at 16.6/34.1/51.3 Hz) and Monkey H’s spectrum on 2020-02-16 (*ap*_off_ = 2.02 logµV^2^, *ap*_exp_ = 1.62, peaks at 21.3/35.3/52.7/66.9 Hz). **(B-E)** Longitudinal recordings of aperiodic features (*ap*_exp_, *ap*_off_) and the aggregate SMR power between 8 – 45 Hz ( *SMR*) (B) are shown alongside lesion sensitivity estimates ⟨*Δ*⟩*_L_* (C) (\**p <* 0.05). Aperiodic shift_Σ_s drove reductions to post-lesion *γ* power (Figure 1E-F) as shown across individual lesions (D). In both monkeys, *SMR* increased after lesions (Monkey U: ⟨*Δ*⟩*_L_* = 2.35, *p <* 1 × 10*^−^*^3^; Monkey H: ⟨*Δ*⟩*_L_* = 1.26, *p* = 0.004) (C) as shown by visualizing the periodic component across individual lesions (E); the gray region indicates the frequency range used to aggregate SMR power in to Σ*SMR*. Note that the power of SMRs was generally stronger in Monkey U (see A as well as bottom of B); this *SMR* amplification likely drove increased post-lesion *β* power in Monkey U (Figure S1B-C), while Monkey H’s *β* power decrease was likely driven by aperiodic shifts. All lesion sensitivity statistics are shown in Table S2.

Meanwhile, lesions increased Σ*SMR* in both monkeys (Monkey U: ⟨*Δ*⟩*_L_* = 2.35, *p <* 1 × 10*^−^*^3^; Monkey H: ⟨*Δ*⟩*_L_* = 1.26, *p* = 0.004). Interestingly, this effect size was larger in Monkey U compared to Monkey H (≈2 vs ≈1 standard deviation). Monkey U also had stronger periodic power content (median Σ*SMR* [IQR]; Monkey U: 40.36 [38.02, 43.12]µV^2^; Monkey H: 23.68 [22.20, 25.56]µV^2^) as visible in Figure 2A-B. Accordingly, periodic *β* power explained more *β* variance in Monkey U than Monkey H (*R*^2^; Monkey U: 0.62; Monkey H: 0.14); this reversed when considering log aperiodic *β* content (*R*^2^; Monkey U: 0.02; Monkey H: 0.66). This difference may also be supported by Monkey H having a flatter neural spectrum (median *ap*_exp_ [IQR]; Monkey U: 1.75 [1.74, 1.78]; Monkey H: 1.64 [1.59, 1.74]). Note that when analyzing individual SMRs, this increase in power was also detected (*SMR_↓_^P^* and *SMR_↑_^P^*); in addition, peak frequencies (*SMR_↓_^f^* and *SMR_↑_^f^*) were reduced in Monkey U while effects on peak widths (*SMR_↓_^W^* and *SMR_↑_^W^*) were mixed (Figure S3C-D, Table S2).

Taken together, spectral parametrization resolved the observed bandpower shifts into two underlying phenomena: aperiodic shifts^22,23,56,57^ and potentiation of Σ*SMR*. These results also show that agreement at the level of bandpower may only exist insofar as a band has a similar aperiodic-periodic constitution across subjects. Here, aperiodic activity originated *γ* power in both monkeys, hence the consistent decrease. However, periodic/aperiodic activity were the main drivers of *β* power in Monkeys U/H respectively, hence the opposing lesion effects. This underlines how critical it is to frame bandpower-centric analyses with spectral parameters. Finally, these effects can also comment on the increase in *T*_rxn_ exclusively observed in Monkey U. Across both monkeys, a moderate correlation existed between *T*_rxn_ and both *α* power (Monkey U: *ρ* = 0.41, *p <* 1 × 10*^−^*^3^, *N* = 100; Monkey H: *ρ* = 0.37, *p* = 0.036, *N* = 65) and *β* power (Monkey U: *ρ* = 0.39, *p <* 1 × 10*^−^*^3^, *N* = 100; Monkey H: *ρ* = 0.33, *p* = 0.051, *N* = 65), replicating past work^87–89^ on a longitudinal scale (all statistics in Table S9 and Table S10). As such, it is reasonable that in Monkey U, where SMRs were amplified to the degree of increasing these bandpowers, *T*_rxn_ also increased; in Monkey H, where SMRs were not constitutive of these bandpowers and were not amplified as strongly, no such increase was observed.

### Spectrum perturbations were movement-specific, spatially diffuse, and temporally unordered

All analyzed data so far has been locked to the initiation of arm reaches (i.e. *T*_rxn_), and it remains unclear whether these perturbations are movement-specific or reflect a global shift in motor cortex function. Our experimental flow did not include a prescribed rest period, but we recomputed spectral features using a window of time immediately before the reach stimulus where the animal was returning their cursor to the center of the screen and awaiting the next trial. While this is not an interpretable rest period, it presents the motor cortex in a different operational mode (performing a predictable movement and holding still at the center). Single-trial arm speeds are shown for both the pretrial and movement periods in Figure S4A. Neither the *γ* (Monkey U: ⟨*Δ*⟩*_L_* = 0.72, *p* = 0.140; Monkey H: ⟨*Δ*⟩*_L_* = −2.42, *p <* 1 × 10*^−^*^3^) nor Σ*SMR* (Monkey U: ⟨*Δ*⟩*_L_* = 1.83, *p <* 1 × 10*^−^*^3^; Monkey H: ⟨*Δ*⟩*_L_* = 0.69, *p* = 0.060) perturbation was replicated in both animals using the pretrial period (Figure S4B, see full statistics in Table S3). Specifically, the strength of the Σ*SMR* increase was attenuated in each animal (Figure S4C) and became insignificant in Monkey H (pretrial vs move; Monkey U: ⟨*Δ*⟩*_L_* = 1.83 vs ⟨*Δ*⟩*_L_* = 2.35; Monkey H: ⟨*Δ*⟩*_L_* = 0.69 vs ⟨*Δ*⟩*_L_* = 1.26). Meanwhile, while Monkey H showed consistent effects on aperiodic parameters and *γ* power, Monkey U was less consistent (Figure S4C). These results indicate that movement may be relevant for demonstrating the spectral effects of covert neuron loss.

Next, we considered whether spectral effects were spatially organized based on proximity to the two electrodes (anode/cathode) for each lesion. Thus far, channels and trials have been aggregated in order to denoise periodogram estimates (Figure S5A); instead, we computed spectral features (e.g. *γ* and Σ*SMR*) for each channel by only aggregating across included trials (Figure S5B). By visualizing the *z* -scored first difference induced by each lesion (denoted *Δ*) on *γ*, the *γ* reduction was diffused across the entire array and wasn’t condensed to the region surrounding the lesion electrodes (Figure S5C). Similarly, Σ*SMR* increased after most lesions in a way that wasn’t spatially focalized (Figure S5D). This was made concrete by computing Spearman correlation between *Δ* for each channel/lesion with its Manhattan distance to the nearest lesion electrode, where electrodes were spaced with a pitch of 400 µm (Figure S6A). There was no spatial correlation for *γ* (Monkey U: *ρ* = 0.00, *p* = 0.997, *N* = 384; Monkey H: *ρ* = −0.05, *p* = 0.360, *N* = 672) and only a weak spatial correlation for Σ*SMR* in Monkey H (Monkey U: *ρ* = −0.01, *p* = 0.921, *N* = 384; Monkey H: *ρ* = 0.13, *p* = 0.004, *N* = 672). Additional spatial correlation statistics for other spectral features are reported in Table S4.

Finally, a unique aspect of our approach is that lesions were repeated within a single animal (Monkey U: *n* = 4; Monkey H: *n* = 7). This invites the possibility that, via mechanisms such as glial remodeling or neural/behavioral compensation, lesion impact could be temporally ordered (i.e. increasing or decreasing with additional lesions). To test this, we considered lesions as repeated measures of the magnitude of *z* -scored first differences (|*Δ*|) for behavioral and spectral features. This totaled 22 features across 4 lesions in Monkey U and 19 features across 7 lesions in Monkey H (Detailed Methods). A Friedman test identified a significant, small difference in |*Δ*| across lesions for Monkey U (*χ*^2^(3) = 9.42, *W* = 0.14, *p* = 0.024), but not for Monkey H (*χ*^2^(6) = 9.89, *W* = 0.09, *p* = 0.130). Corrected, pairwise Wilcoxon signed-rank tests (two-tailed) found that effects decreased from Lesion 1 to both of Lesions 2/3 in Monkey U (1 vs 2: *T* = 34, 72.7 % decreased, *p* = 0.024; 1 vs 3: *T* = 32, 68.2 % decreased, *p* = 0.024; Figure S6B). This suggested the possibility of a decreasing trend in effects with repeated lesions, however Page’s trend test for this directional hypothesis failed to reject the null hypothesis in both animals (Monkey U: *L* = 561.0, *p* = 0.220; Monkey H: *L* = 2096.0, *p* = 0.742). Similarly, by computing Spearman correlation between |*Δ*| and the temporal position of the corresponding lesion (i.e. 1, 2, *. . .*), there was not a significant correlation (Monkey U: *ρ* = −0.06, *p* = 0.600, *N* = 88; Monkey H: *ρ* = 0.10, *p* = 0.260, *N* = 133). This suggests that while repeated lesions were heterogeneous, there was no temporal ordering of effect sizes.

### Spectrum perturbations outlasted behavioral effects in a state space model

So far, lesion impact has been assessed by permutation tests on a univariate lesion sensitivity statistic ⟨*Δ*⟩*_L_*. But, this approach does not estimate the duration of lesion impact, only the presence of a next-day effect. To strengthen these results, we set up a LG-SSM^66,90^ (Figure 3A). This decomposed observed time-series (*y* in Figure 3A) into a component arising from a linear dynamical system (i.e. from *x*), a component driven by inputs (i.e. from *u*), and noise. By treating the lesions as inputs, this model discerned the duration of time for which lesions had a significant input on the time-series that could not be explained by underlying temporal dynamics. This confirmed and further differentiated the sensitivity of spectral and behavioral features to neuron loss. We first fit the model to Behavior (*T*_rxn_, *v*_reach_, *θ*_err_) and Periodogram (*ap*_exp_, *ap*_off_, Σ*SMR*, *δ*, *θ*, *α*, *β*, *γ*) datasets (example reconstructions are shown in Figure S7A). Permutation of lesion days tested the significance of the 5 day lesion input for each feature (*d*_1_, *d*_2_, *. . .*, *d*_5_), and determined whether structured lesion inputs improved marginal log likelihood (SSM_LL_). As expected, lesion inputs significantly improved SSM_LL_ for all datasets except for Behavior in Monkey H (top of Figure 3B-C, see likelihoods and *p*-values in Table S5). Accordingly, the only significant behavioral input was a 1 d increase to *T*_rxn_ in Monkey U (*d*_1_ = 1.92, *p* = 0.030) (bottom of Figure 3B). Meanwhile, the model estimated a multiday reduction to *γ* power (duration; Monkey U: 2 d; Monkey H: 3 d), with effects on the order of a standard deviation for three days (Monkey U: *d*_1_ = −1.89, *p* = 0.020, *d*_2_ = −1.25, *p* = 0.040, *d*_3_ = −1.17, *p* = 0.053; Monkey H: *d*_1_ = −1.28, *p* = 0.008, *d*_2_ = −1.23, *p* = 0.008, *d*_3_ = −0.85, *p* = 0.020) (bottom There was also an increase to Σ*SMR* (Monkey U: 2 d; Monkey H: 1 d) with effects of a similar magnitude (Monkey U: *d*_1_ = 1.20, *p* = 0.020, *d*_2_ = 1.28, *p* = 0.020; Monkey: *d*_1_ = 0.95, *p* = 0.011) (see all parameters in Table S6, Table S7). Additional behavioral and spectral features can be seen in Figure S7B-C. Overall, spectral effects outlasted behavioral effects for each monkey (i.e. elevated *T*_rxn_ for 1 d in Monkey U), suggesting a stronger influence of neuron loss on the spectrum, relative to behavior, in this covert context.

**Figure 3:**
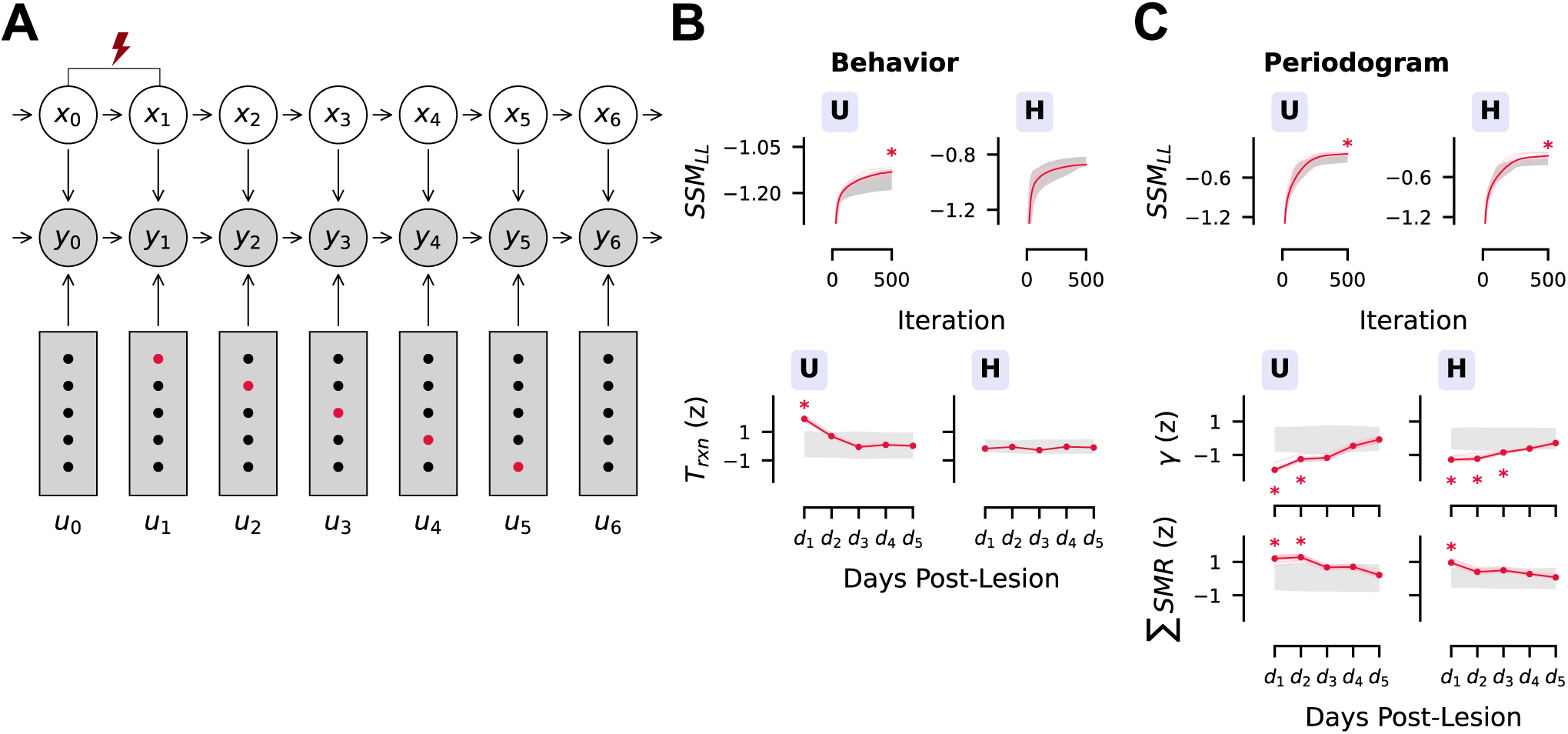
Spectrum perturbations outlasted behavioral effects. **(A)** A linear Gaussian state space model (LG-SSM) was used as an alternative framework to quantify the impact of lesions; this approach also quantifieda temporal extent of lesion impact. Specifically, the model represented lesions as 5 day external inputs (*u_t_*) on observed features (*y_t_*) that couldn’t be accounted for by the linear dynamics of a low-dimensional state (*x_t_*) (see Detailed Methods). Lesion inputs were a vector of 5 values, where the value at index *i* was 1 if it was *i* days after a lesion, and 0 otherwise; this is shown by the crimson and black dots. By fitting this model to collections of *z* -scored time series, this allowed estimation of lesion impact across the 5 day, post-lesion period. For each feature, this was measured by *d*_1_, *d*_2_, *. . .*, *d*_5_ (not shown), the standardized input strength of lesions on each post-lesion day. Note that this graphical model captures only a small segment of the longitudinal time series which contained multiple lesions. **(B-C)** On the Behavior (B) and Periodogram (C) datasets, permutation tests determined the significance of SSM_LL_ (top) and *d_i_* (bottom) (\**p <* 0.05). Gray shading indicates 95 % intervals for SSM_LL_ fitting curves (top) or input estimates (bottom) across 1000 permuted null models. Lesion inputs increased SSM_LL_ for the Behavior dataset in Monkey U (B, top) and the Periodogram dataset (C, top) in both monkeys. This approach confirmed the only detectable behavioral effect of le_Σ_sions, a 1 d increase to *T*_rxn_ in Monkey U (B, bottom). It also confirmed the reduction to *γ* power and increase to *SMR* (C, bottom). These spectral effects o_Σ_utlasted any behavioral impact (Monkey U: Behavior=1 d, *γ*=2 d, *SMR* =2 d; Monkey H: Behavior=0 d, *γ*=3 d, *SMR* =1 d), suggesting that the spectrum is more sensitive than behavioral metrics to neuron loss. As visible on the bottom of (B) and (C), effect sizes were on the order of a standard deviation. For full LG-SSM statistics, see Table S5, Table S6, Table S7, and Table S8; additional visualizations are in Figure S7.

### Time-frequency signatures of lesions were personalized

The analysis so far has used static descriptions of spectral activity during arm reaching, but it is known that *α* and *β* activity is modulated while preparing,^91–93^ initiating,^83,94,95^ and carrying out^96,97^ motor commands, and that these time-frequency dynamics are perturbed in symptomatic stroke.^1,11–16^ As such, a non-negative time-frequency tensor was formed with squared wavelet coefficients across days (Figure 4A, left; see Detailed Methods). This is a high-dimensional, multiway data structure; to generate a parsimonious representation that is useful scientifically, one can use domain knowledge to extract classical features (e.g. bandpower) or use a data-driven parametrization (e.g. spectral parametrization). Here, classical features measure the movement-related dynamics of *β* activity which is known to desynchronize at movement onset (MRBD) and re-synchronize at movement cessation (PMBR).^85,86^ These variables (MRBD, PMBR) were computed by normalizing wavelet power around median *SMR^f^* in each animal (visualized as *β*_tf_ in Figure S8A) relative to a baseline (*β*_ref_) (Figure S8B). The diverging impact on *β* bandpower observed previously also appeared in *β*_ref_ (Monkey U: ⟨*Δ*⟩*_L_* = 1.18, *p* = 0.015; Monkey H: ⟨*Δ*⟩*_L_* = −2.11, *p <* 1 × 10*^−^*^3^). This effectively increased the dynamic range for MRBD/PMBR activity in Monkey U while shrinking this range in Monkey H; accordingly, MRBD was augmented (i.e. larger decrease) in Monkey U (⟨*Δ*⟩*_L_* = −1.31, *p* = 0.005) but attenuated (i.e. smaller decrease) in Monkey H (⟨*Δ*⟩*_L_* = 0.958, *p* = 0.019) (Figure S8C, Table S2). Meanwhile, PMBR decreased in Monkey U (⟨*Δ*⟩*_L_* = −1.46, *p* = 0.005) but was unaffected in Monkey H (⟨*Δ*⟩*_L_* = 0.641, *p* = 0.086). This agrees with literature showing heterogeneous measurements of MRBD/PMBR changes in stroke cohorts.^13,15,16^

**Figure 4:**
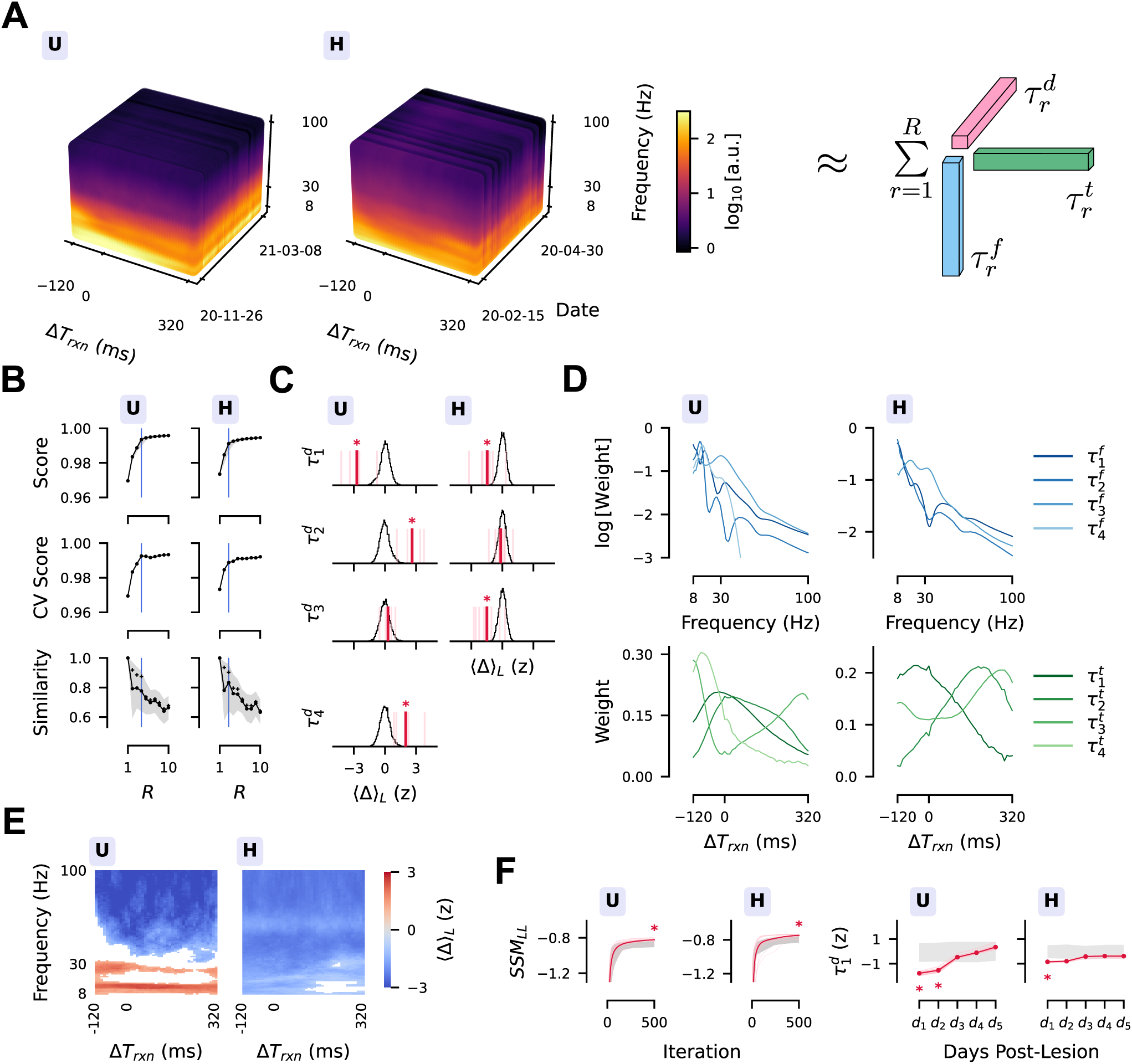
Signatures of lesions in time-frequency dynamics were personalized. **(A)** Squared wavelet coefficients yielded a time-frequency tensor across days. An *R*-rank approximation was found using the CPD, identifying constituent factors across days (*τ_r_^d^*), frequency (*τ_r_ ^f^*) and time (*τ_r_ ^t^*). **(B)** Based on high reconstruction score, increasing cross-validation score, and high model similarity across initializations, *R* = 4 and *R* = 3 were chosen for Monkeys U and H. Lines are the mean with shaded 95 % percentile intervals. Reconstructions scores were computed across 1000 initializations for each rank, while 50 cross-validation scores were computed from 25 instances of 2-fold cross validation. Similarity scores captured the 999 similarities (Detailed Methods) with the highest-scoring initialization; “+” markers isolate the mean similarity of the 99 next-best models. Note that this metric drops off meaningfully after the selected ranks in each monkey. **(C-D)** Both *τ*_1_ and *τ*_2_ had broad, unimodal temporal weights alongside broad frequency weights, although *τ*_1_ had increased relative weight on *γ* frequencies (D). In both monkeys, *τ_1_^d^* was reduced by lesions (Monkey U: ⟨*Δ*⟩*_L_* = −2.71, *p <* 1 × 10*^−^*^3^; Monkey H: ⟨*Δ*⟩*_L_* = −1.47, *p <* 1 × 10*^−^*^3^), echoing the impact on *γ* bandpower previously observed, while *τ_2_^d^* was preserved in Monkey H (⟨*Δ*⟩*_L_* = −0.15, *p* = 0.668) and increased in Monkey U (⟨*Δ*⟩*_L_* = 2.62, *p <* 1 × 10*^−^*^3^) (C) (\**p <* 0.05). Meanwhile, *τ*_3_ had bimodal frequency and temporal weights, reflecting movement-related *β* modulations, while *τ*_4_ was unique to Monkey U and captured pre-movement *α* and *β* power. Effects on these factors revealed a divergent lesion impact on SMR dynamics, mirroring the analysis of *β*_ref_/MRBD/PMBR in Figure S8. **(E)** The lesion sensitivity of individual wavelet coefficients, shown here, was demixed by the CPD (mask indicates significance at *p <* 0.05). **(F)** State space modeling affirmed that lesion inputs improved SSM_LL_ of the *τ_r_^d^* (Table S5) and revealed a 2 d reduction to *τ*_1_*^d^* in Monkey U and a 1 d reduction in Monkey H (\**p <* 0.05) (Table S8).

However, these perturbations may be driven by lesion impacts that are temporally dynamic/static or focused on a wide/narrow range of frequencies. To make this distinction, dimensionality reduction techniques can reduce high-dimensional data into constitutive factors^98^. For our longitudinal time-frequency tensor, the nonnegative CPD^62–64^ was used to approximate the tensor as a sum of *R* rank 1 component tensors (Figure 4A). The CPD has been used in both electroencephalography studies^61,99,100^ and invasive population recordings.^101,102^ Simulation work has also shown it to be a suitable model for LFP recordings,^103^ but use has been limited, especially for longitudinal datasets.^104,105^ Importantly, the CPD has strong uniqueness properties, contrasting approaches like principal component analysis which impose constraints on uncovered factors such as orthogonality.^101,106^ If this unique solution can be discovered, then the decomposition factors can be directly interpreted.

Model selection metrics, such as high model similarity across initializations (Detailed Methods, Figure 4B, Figure S9A) suggested ranks of *R* = 4 for Monkey U and *R* = 3 for Monkey H. The estimated time-frequency factors (*τ_r_ ^f^* and *τ_r_^t^*) are shown in Figure 4D alongside the lesion sensitivity of date factors *τ_r_^d^* (Figure 4C, see *τ_r_^d^* trends in Figure S9A). In both monkeys, *τ*_1_ *^f^* and *τ*_2_ *^f^* had broad frequency content with *τ_1_ ^f^* showing stronger relative frequency weight on the *γ* range. Both temporal factors (*τ*_1_ *^t^* and *τ*_2_*^t^*) had broad, unimodal weights. In addition, *τ*_1_*^d^* was reduced by lesions (Monkey U: ⟨*Δ*⟩*_L_* = −2.71, *p <* 1 ×10*^−^*^3^; Monkey H: ⟨*Δ*⟩*_L_* = −1.47, *p <* 1 ×10*^−^*^3^) while *τ_2_^d^* was preserved in Monkey H (⟨*Δ*⟩*_L_* = −0.15, *p* = 0.668) and increased in Monkey U (⟨*Δ*⟩*_L_* = 2.62, *p <* 1 × 10*^−^*^3^) (Figure 4C). These factors recapitulated results from the static spectrum analysis, where *τ*_1_ captured a broad power reduction (especially in *γ* frequencies), while *τ*_2_ reflected the relative preservation of activity in *α* and *β* bands from increased Σ*SMR*. *Τ_2_^d^* also had a moderate correlation with *T*_rxn_ (Monkey U: *ρ* = 0.42, *p <* 1×10*^−^*^3^, *N* = 100; Monkey H: *ρ* = 0.37, *p* = 0.008, *N* = 65) (Table S9, Table S10) just like *α*/*β* power. Meanwhile, *τ*_3_ captured movement-related modulations of SMRs featuring a bimodal *τ_3_ ^f^* alongside a bimodal *τ_3_^t^* with a similar temporal trajectory as empirically computed for *β*_tf_ in Figure S8A. These movement-related dynamics were thus demixed from the more static factors (*τ*_1_ and *τ*_2_) (Figure 4D). In Monkey U, this factor was also paired with *τ*_4_ which had a sharp temporal activation before movement onset and concentrated frequency content in the *α* and *β* range. These factors re-iterated the personalized signature of lesions in *β*_tf_ activity. For Monkey U, *τ_4_^d^* was elevated (⟨*Δ*⟩*_L_* = 2.01, *p <* 1 × 10*^−^*^3^) while *τ_3_^d^* was preserved (⟨*Δ*⟩*_L_* = 0.29, *p* = 0.522), compatible with elevated *β*_ref_ increasing the MRBD dynamic range. In Monkey H, *τ_3_^d^* was reduced (⟨*Δ*⟩*_L_* −1.49, *p <* 1 × 10*^−^*^3^), compatible with a generic shrinking of MRBD dynamic range (Figure 4D). Finally, an LG-SSM model was also fit to the *τ_r_^d^* ; structured lesion inputs improved the model (left of Figure 4F, Table S5) and featured a 2 d post-lesion reduction to *τ_1_^d^* in Monkey U alongside a 1 d reduction in Monkey H (right of Figure 4F) while confirming the lesion impact on other factors (full statistics in Table S8). Overall, tensor decomposition demixed the lesion effects on individual wavelet coefficients (Figure 4E), unifying static spectrum impact with the personalized effect on MRBD/PMBR metrics.

## Discussion

This study measured the previously unexplored effects of covert neuron loss in LFP data as a foundation for clarifying the causal relationships between functional motor impairment and spectrum shifts after stroke. Covert neuron loss–which preserved success rates and minimally impacted reaching behavior–was repeated within two NHPs. Certain spectral signatures of symptomatic stroke were recapitulated, while still more effects were suggested, despite the preservation of behavioral ability. For instance, just as in symptomatic stroke, a pronounced reduction in *γ* power was observed alongside other bandpower perturbations.^4,19–22^ In alignment with modern analyses of the post-stroke spectrum,^22,23,56,57^ this effect was driven by aperiodic shifts.^55^ Spectral parametrization also revealed that the power of *α*/*β* SMRs (Σ*SMR*) increased after lesions. Additional analyses determined that these spectral perturbations were accentuated during movement-related epochs, spatially diffused across the channels of the Utah array, and did not show an ordering of effect size with repeated lesions. These findings contrasta recent study that observed a reduced low *γ* rhythm at 7 days post-ischemia in mice,^22^ although a direct comparison is challenging as recordings were made under general anesthesia. A more similar study analyzed the post-injury movement-related spectrum,^23^ but found that peak power in the 15 – 16 Hz range was unperturbed. This discrepancy may be driven by imperfect identifiability of spectral parametrization,^107^ differences in rodent and primate neurophysiology, or may reflect a difference in covert and symptomatic spectral responses that could have functional significance.

This latter lens is what relates these results to clinical interest in covert stroke^35,38^ as well as to theoretical work modeling the recovery boundary, a threshold of neuron loss beyond which a cortical network cannot rapidly compensate and maintain its function.^34^ In past stroke research, measurements have been made almost exclusively beyond this boundary, where spectrum shifts coincide with behavioral breakdown. This has produced correlations between features such as the *ap*_exp_^23,56,57^ and the *δ*-*α* ratio^5,7^ with behavioral recovery. Here, we show that the spectrum is perhaps more sensitive to neuron loss than previously understood, and that spectrum perturbations can be induced by covert injury independent of substantial behavioral deficit. In fact, the only behavioral consequence of the lesions was a 1 day increase to *T*_rxn_ in Monkey U ranging from ∼14 – 17 ms across lesion sensitivity and state space modeling statistics (see Table S2 and Table S6). This coheres with previous reports that covert neurological symptoms can act as indicators for covert stroke,^108^ although these measurements show that spectral features, which had pronounced, multi-day signatures of covert neuron loss, may be more effective. This work thus adds more causal context to previously observed correlations,^3,23^ and while these results are exploratory and don’t definitively defend a causal structure, they motivate further work leveraging controlled neuron loss across various scales. As a reference, reaching deficits have been recorded across a 5-week timescale after complete aspiration of the arm motor cortex,^109^ suggesting some intermediate injury severity^34^ at which acute, next-day symptoms should emerge. Pursuing larger lesions with increased current and/or duration is an important next step in order to induce more substantial behavioral breakdown, which might be articulated by the disappearance, emergence, or intensification of certain LFP signatures. For instance, recent work has proposed that restoring disrupted movement-related *δ* oscillations after stroke may be a useful therapeutic target,^17,18,110^ although such a disruption was not identified in this dataset. In addition, the observed increase of Σ*SMR* may signify covert compensation and become undetectable at the clinical boundary, as suggested by recent work.^23^

One possible interpretation for aspects of the lesion response is an increase in local cortical inhibition, shifting excitation-inhibition (E:I) balance. Slowing and magnification of *β* rhythms has been measured upon treatment with diazepam, a GABAergic agonist that increases the amplitude of inhibitory post-synaptic potentials.^111–113^ Moreover, *ap*_exp_ has been shown to steepen with increased inhibitory conductance in simulation, increased I:E synaptic density balance, and upon delivery of anesthesia in monkeys.^114^ All of these inhibitory signatures were represented in Monkey U’s response, while elements were also present (such as SMR magnification) in Monkey H. Importantly, this interpretation bridges the current study with observations of increased local inhibition after stroke,^22,115–119^ although most prior work has emphasized how elevated tonic inhibition–related to extrasynaptic GABA*_A_* receptors and the concentration of ambient extracellular GABA–impedes behavioral recovery,^22,115–118^ while the aforementioned LFP features are more closely tied to synaptic, phasic inhibition.^119,120^ This suggests that mechanisms such as insertion of synaptic GABA*_A_* receptors may be activated following covert neuron loss to mitigate excitotoxic neuron loss,^119,121^ but this cannot be confirmed without additional data.^115,119,122^ The observed responses may also reflect local neuroinflammation as there is emerging evidence that both microglia^123^ and astrocyte^124^ recruitment can alter neural network oscillations. Untangling the interplay between phasic and tonic inhibition, as well as neuroinflammation, in post-injury cortical networks may be crucial for advancing stroke treatments.

Complementing the analysis of static, peri-movement spectra, the exploration of movement-related time-frequency dynamics revealed diverging effects on baseline, pre-movement *β* activity (*β*_ref_) in the two animals, leading to increased dynamic range and exaggerated MRBD in Monkey U and the opposite effect in Monkey H. This was paired with a novel application of the CPD–an interpretable tensor decomposition–which recapitulated aspects of the static spectral signatures described above while affirming this divergent effect on movement-related *β* dynamics. Interestingly, this heterogeneity is also consistent with the sparse literature examining movement-related *β* modulations after stroke. For example, a recent study showed that patients starting with strong *α* desynchronization further increased this desynchronization during rehabilitation (like Monkey U), while weak desynchronization was further decreased (as in Monkey H).^16^ Other studies have failed to identify a cohort-level effect on MRBD^15^ or have reported an overall reduction.^13^ These results signify that part of the response to neuron loss may be personalized, where consequences are dictated by bio-behavioral statistics. For instance, Monkey H was 14 and near end of life while Monkey U was 11 and in good health at the time of recording. Age^125,126^ and general health^125,127^ has been associated with worsened functional and biological consequences of ischemia (although other measurements challenge these correlations^128^); moreover, age is known to erode endogenous inhibitory tone^129^ while shifting the neural spectrum’s aperiodic-periodic constitution by flattening *ap*_exp_ and reducing Σ*SMR*,^55,130^ as can be seen when comparing Monkeys U/H in Figure 2A-B. Additional data will be critical for dissecting the relationships between age/health, spectral statistics, and the neural-behavioral response to neuron loss.

Two methodological considerations were important to this work. First, spectral parametrization,^55^ state space modeling^90^, and tensor decomposition^64^ all added interpretive depth to the measured impact of neuron loss, complementing the information extracted from classical LFP features. Spectral parametrization improved interpretability of estimated bandpower effects by decoupling an enhancement in periodic rhythms from a shift in aperiodic structure. State space modeling leveraged the time-series structure of the data to estimate the duration of lesion inputs and to ensure that these effects did not arise from underlying temporal dynamics. Meanwhile, tensor decomposition is an emerging framework for visualizing and analyzing neuroscience tensors,^101–103^ and was used to find interpretable time-frequency dynamics affected by lesions. While spectral parametrization approaches are becoming clinically relevant,^57^ the state space modeling and tensor decomposition approaches were tailored to our dataset in order to answer specific scientific questions. We advocate for their use in datasets with similar properties (longitudinal designs with known intervention points), but it is unlikely that these techniques can be immediately implemented in the clinic. A second methodological consideration was the electrolytic lesion platform.^53^ The platform permits precise, mm-level control of lesion extent by tuning electrical current and duration; this facilitated calibration to an approximate lesion volume of ∼1 mm^3^.^53^ Moreover, this lesioning platform is neuroelectrophysiology compatible; it induced neuron loss from an already implanted array, permitting immediate pre-/post-lesion recordings during behavior. Such recordings are more difficult in contemporary approaches to neuronal termination such as endovascular occlusion^131^ or photothrombosis,^132^ both of which require a surgery that can disrupt an already-implanted neuroelectrophysiology device. In a similar vein, this platforms permits repeated lesions (that do not cause array degradation^54^) without additional surgeries. Collectively, these features permitted recording of longitudinal, baseline-referenced responses to covert neuron loss. In addition, the platform enabled a within-subject experimental design using permutation testing, factoring out inter-individual heterogeneity and enabling sensitive detection of neuron loss impact.

A first limitation is that, although inter-animal variability in age or health will become a strength of our paradigm with more subjects, the study’s broader conclusions are limited without going beyond the 2-monkey convention of primate systems neuroscience. However, we used strict behavioral filtering (Detailed Methods) to include well-stereotyped behavioral conditions in each animal (e.g. unaffected by Monkey H’s visual deficit). Moreover, our statistical approach used each monkey as their own empirical control, ensuring that lesion impacts reflected genuine departures from expected day-to-day, animal-specific spectral/behavioral fluctuations. A second limitation is that 2D endpoint position in a simple reaching behavior may not be adequate to reveal subtle lesion effects. While this behavior is standard in motor systems neuroscience, and was analyzed from multiple analytic perspectives, coupling this lesion model with higher-dimensional reach/grasp representations^133^ or ethological models^134^ of naturalistic behavior^135–137^ may unify brain-behavior responses to neuron loss. A third limitation is that while these results point to the scientific potential of the lesions for studying covert and symptomatic stroke, the analogy to ischemic injury is incomplete. For one, ∼90 % of covert strokes are subcortical.^35^ However, the lesion platform is compatible with linear microelectrode arrays as shown in our previous work^53^. These arrays permit subcortical lesions, contrasting the capacity of optical approaches like photothrombosis or the consequences of distal middle cerebral artery occlusion. Subcortical lesions are an attractive direction for future work, especially due to our expanding perspective on the role subcortical circuits play in expressing innate and learned motor skills, including the arm reaches studied in our paradigm.^138^ Secondly, epidemiological studies of covert stroke have found that advanced age^38^ and hypertension^35,37,38,41,139^ are the largest risk factors, while the monkeys lesioned here were middle-aged (11 – 14 years) without known cardiovascular dysfunction. This mismatch encourages the replication of our results in older, hypertensive^140^ macaques. Thirdly, electrolytic lesions are not vascular injuries, despite inducing a similar consequence (neuron loss). Our model does not capture the vascular origin and molecular/cellular pathology of covert stroke. However, the motivating principle of our model views neuron loss as the core functional consequence of stroke, wherein functional recovery takes place alongside reorganization^26–28,31,141^ or potentiation^29^ of the remaining neuronal population. Our model allows us to track changes in the LFP in relation to neuron loss, a relation that we believe should exist regardless of pathophysiological mechanism. This is supported by recent work showing that ministrokes of cerebral cortex decoupled cerebral blood flow and LFP, and found that the LFP more robustly indicated the functional consequences of neuron loss beyond the subacute phase.^142^ As such, the LFP describes functional outcomes moreso than acute vascular dysfunction. Our results, and their integration with the spectral impact of symptomatic stroke, further support this. Overall, the decoupling of neuron loss from vascular pathology is an intentional aspect of this model, which complements other animal models of stroke that use a vascular mechanism.^142–144^ A future opportunity for animal model revision is the unification of the vascular-origin infarcts with chronic longitudinal neuroelectrophysiology. A final limitation is that, although this work focused on LFP, it is at the resolution of neuronal population spiking activity where computations related to motor planning and execution appear to occur.^145^ Already, population statistics are shedding light on post-stroke reorganization;^110,146^ fortunately, this data is also recorded from the implanted array, and it will be imperative to explore spiking-level lesion signatures in future work.

## Author Contributions

KM was responsible for conceptualization, data curation, formal analysis, investigation, methodology, software, validation, visualization, writing, and review/editing. SEC was responsible for conceptualization, data curation, investigation, methodology, supervision, and review/editing. PN was responsible for conceptualization, funding acquisition, project administration, resources, supervision, and review/editing. The members of the Brain Interfacing Laboratory provided additional support in review/editing.

## Acknowledgments

We thank Iliana E Bray for their effort in collecting core experimental datasets. We thank Stephen I Ryu for nonhuman primate array implantation. We thank Mackenzie Risch, Kristina Lebedev, and Michelle S Wechsler for expert surgical assistance and veterinary care. We thank Sam Baker for veterinary support and Kimberly Chin for administrative support.

