## Supplementary Material for "Signatures of covert neuron loss in the local field potential of motor cortex"

### Detailed Methods

#### Animal care

Two adult male rhesus macaques (*Macaca mulatta*), ages 11 (Monkey U) and 14 (Monkey H) at the start of recording, were implanted with 96-channel Utah arrays using stereotaxic coordinates in the contralateral motor cortices (Blackrock Microsystems, Salt Lake City, UT, USA). Utah arrays with 1.5 mm iridium oxide electrode shanks were implanted in Monkey U, and with 1.0 mm platinum electrode shanks in Monkey H. Monkey U was implanted in 2017 with three arrays in the left medial/lateral primary motor cortex, as well as left premotor cortex. Monkey U performed reaches with his preferred right arm. Monkey H was implanted in 2012 with two arrays in the right primary motor cortex and premotor cortex and performed reaches with his preferred left arm. All animal procedures were reviewed and approved by Stanford University's Institutional Animal Care and Use Committee.

#### Behavior

During simultaneous microelectrode recordings of the local field potential (LFP) from primary motor cortex (M1) (Figure 1B, left), two monkeys (U and H) performed a traditional center-out reaching task nearly every day, at approximate 24-hour intervals, for months. Monkey U performed this task on 102/103 days spanning 2020-11-26 to 2021-03-08 (no recording on 2021-01-25). Monkey H performed the task on 73/76 days spanning 2020-02-15 to 2020-04-30 (no recording on 2020-03-21, 2020-03-22, and 2020-04-23). In addition, Monkey H chose not to perform at least 100 trials on 2020-02-29, 2020-03-01 and 2020-04-03; these days were discarded, yielding a dataset of 70 days.

Monkey U played with 8 equally spaced radial targets, while Monkey H had a visual deficit and played with the bottom and bottom left targets removed, leaving 6 target locations. In addition, Monkey H showed poor stereotyped reaching toward the top and top right targets (see examples of delayed, highly-variable movement onset times in Figure S1A). Since this study used descriptive daily statistics of reaching ability and refrained from condition-specific analysis, these reaches were discarded in order to isolate engaged, consistent movements across days; this left 4 targets for analysis. However, the task described below was delivered using the original pool of 6 targets.

Each trial started with a green target (12 mm diameter) appearing at the origin of the workspace (0,0). The monkey would move their arm to hover the cursor on the target, and after holding within the target for 300 ms (Monkey H) or 200 ms (Monkey U), one of eight (Monkey U) or 6 (Monkey H) radial targets would appear at a distance of 100 mm (Monkey U) or 60 mm (Monkey H) from the origin. The target selection was pseudo-random to ensure even trial counts while preventing predictions of trial location. The monkey would then move their arm to the radial target, and juice would be dispensed if the target was held for the aforementioned durations. A trial would fail if the target was not acquired within 2 s.

To track reach kinematics, a reflective bead was secured to the third and fourth digits of the hand and optically tracked at 60 Hz with an infrared camera (Polaris Spectra, Northern Digital Inc., Waterloo, Canada). The bead's position was projected onto a computer screen as a task cursor, aligned such that natural, straight extension of the arm mapped to the origin (0, 0). The task also enforced a maximum permitted distance from the screen, requiring the monkeys to reach fully outward and make whole-arm movements within a defined work space of approximately 0.25 $m^3$. All behavioral data streams and control signals for the infrared camera, experimental rig lights, juicer system, and the behavioral task were collected, synchronized, and controlled at 1 kHz using LiCoRICE, an open-source platform for realtime processing.[1]

For each successful trial, these kinematics were converted to a reaction time ($T_{\text{rxn}}$), reach speed ($v_{\text{reach}}$) and angular error ($\theta_{\text{err}}$) (Figure 1B). To do so, kinematic positions ($p_x(t)$, $p_y(t)$), were converted to smoothed $v_x(t)$ and $v_y(t)$ velocity signals in units of cm $s^{-1}$ using a 2nd order, 100-ms Savitsky-Golay filter. The scalar speed was computed for each time point by $s(t) = \sqrt{v_x(t)^2 + v_y(t)^2}$. Next, peaks in speed were detected using `scipy.signal.find_peaks`,[2] with a minimum peak height of 1 cm s$^{-1}$ and a minimum peak width of 20 ms. The first peak that was within 80 % of the largest peak was selected as the initial ballistic reach to the target. The height of this peak, in cm s$^{-1}$, defined the metric $v_{\text{reach}}$. The last upward crossing of $0.15 \times v_{\text{reach}}$ preceding the peak was $T_{\text{rxn}}$. Finally, for the 100 ms after summiting $v_{\text{reach}}$, the average cosine distance between the velocity vector and the vector between current position and the target defined $\theta_{\text{err}}$.

To ensure behavior was stereotyped, this behavioral identification had to pass numerous filters, listed below. In general, these filters would reject the majority of Monkey H reaches toward the top and top right targets, hence their exclusion. An example day of accepted and rejected trials is shown in Figure S1A. The percentage of successful trials admitted each day is shown in the top panel of Figure S1B as the metric rate$_{\text{acc}}$. Importantly, these filters were also not significantly over- or under-activated by lesions (top of Figure S1C; see Statistics section in this document). The filters were:

1. $150 \leq T_{\text{rxn}} \leq 750$ ms, ensuring the animal wasn't guessing the target location and was paying attention
2. In the 50 – 250 ms period before $T_{\text{rxn}}$ the arm had to be stationary ($s(t) <$10 cm s$^{-1}$), ensuring comparable starting arm posture for all epochs
3. $\theta_{\text{err}} < 90°$ to ensure movement was directed toward quadrant of presented target
4. $v_{\text{reach}}$ achieved within 350 ms of $T_{\text{rxn}}$, ensuring $T_{\text{rxn}}$ rapidly transitioned to a ballistic reach

Up to 750 of these stereotyped trials were included in the dataset each day. Epochs were extracted spanning 150 ms before to 350 ms after $T_{\text{rxn}}$. Note that by the behavioral filters, it was assured that the 150 ms period before $T_{\text{rxn}}$ started with a stationary arm, and that the 350 ms after $T_{\text{rxn}}$ featured the attainment of $v_{\text{reach}}$. This helped ensure that analyzed neural activity extracted from these epochs unfolds alongside consistent movements.

On each recording day, these trials were initially presented in a session of up 750 successful reaches, depending on voluntary engagement. Next, a sham/lesion break was taken, before a second session would continue for as long as the animal chose to participate. Trials from these second sessions were not analyzed for Monkey U, but were included where necessary on non-lesion days in Monkey H to reach up to 750 trials per day (due to variable engagement and aforementioned target exclusions). In total, the dataset spanned 102/103 consecutive days for Monkey U (median [IQR] trials per day: 728 [718, 735] trials; total: 74,054 trials) and 70/76 days for Monkey H (619 [317, 750] trials; 36,827 trials). On each day, the medians of $T_{\text{rxn}}$, $v_{\text{reach}}$, and $\theta_{\text{err}}$ were used for subsequent analysis.

#### Electrolytic lesions

Throughout this period of time, small electrolytic lesions were delivered to the motor cortex using a previously reported electrolytic lesioning platform.[3,4] In brief, a circuit was designed to deliver precise, controlled amounts of current (to a precision of 10 µA) using a pair of adjacent electrodes as the anode/cathode respectively to pass DC current through tissue. This approach can cause neuron loss via tissue heating,[5,6] electroporation,[7] and local changes in pH[5]. By selecting the amplitude and duration of the current, the lesion magnitude can be controlled. Through calibration using using histology in *ex vivo* ovine and porcine brains and *in vivo* anesthetized porcine and macaque brains,[3] a parameter range of 150 – 170 µA for 30 – 45 s was identified as being suitable for evoking small, $\approx$1 $mm^3$ lesions to the macaque motor cortex. This lesion volume is a fraction of $\sim$20 $mm^3$ photothrombotic[8] or $\sim$100 $mm^3$ aspiration[9] lesions previously evoked in macaque sensorimotor cortex. This range was explored within these experiments and Monkey U received four 30 s lesions, while Monkey H received ten lesions mostly at a duration of 45 s; full parameters are available in Table S1. These variations stayed within the prescribed range while providing additional information for further calibration of lesion parameters. The only exceptions were Monkey H's third lesion which was at 130 µA for 50 s and Monkey H's final tenth lesion, which was used as an opportunity to explore a larger current (480 µA for 30 s). This study focused on the last 7 lesions for Monkey H, as the first lesion had recording issues while Monkey H refused to participate after the third lesion. Hence, only the last 7 lesions spanned a duration of time where daily recordings were consistent, a feature vital to our statistical approach (see Statistics).

To integrate the lesions into the experiment, as previously mentioned, each recording day started with an initial session of up to 750 successful trials. A sham-/lesion- break would then be taken, typically lasting around 15 minutes. The sham breaks were included to prevent the animal from differentiating the experimental flow of lesion- and non-lesion-days. On lesion days, the circuit would be used to painlessly induce neuron loss during the break. Importantly, this helped ensure animals were blinded to the occurrence of lesions. After each lesion, a medical doctor with expertise in handling primates performed a neurological and physical examination (to the extent possible while the animal was seated in the chair) to evaluate pupillary light response, skin color/turgor, and bilateral arm muscle strength. After most analyzed lesions, ear temperature was taken while for all included lesions, a blood sample was taken from ear capillaries to measure lactate levels alongside other gases (CG4+ cartridge, iStat, Abbott Labs, Chicago, Illinois, USA). In addition, extensive post-lesion monitoring was conducted by members of the Brain Interfacing Lab alongside veterinarians in the Department of Comparative Medicine and animal care staff at Stanford University. These observations were conducted in-person and over remote camera feeds (DNZ30TL2R, Teledyne FLIR, Thousand Oaks, CA) to make detailed records of in-cage behavior (mobility, usage of contralesional arm, posture), sleep, urine/feces production, fluid intake, and appetite. Observation continued for a minimum of 6 h after lesions and often continued for much longer until $\approx$1AM, before resuming as early as 4AM the following day. No abnormalities were observed after any analyzed lesion except for occasional mild restlessness during sleep for Monkey H. Post-lesion medical assessments did not show any neurological impairment, and contralesional arm muscle strength was always graded at 4+ or 5. Blood tests and temperature measurements did not indicate any abnormalities for any analyzed lesion. For complete descriptions of temperature, blood gases, and arm strength, see Table S1.

#### In vivo electrophysiology

As mentioned previously, both Monkey H and Monkey U were intracortically implanted with Utah microelectrode arrays (Blackrock Neurotech, Salt Lake City, UT, USA) in motor regions, including the arm region of M1. During experimental sessions, neuronal data was collected at 30 kHz from M1 and transmitted from a BlackRock CerePlex E headstage to a Blackrock Neural Signal Processor system over fiber optic cable (Blackrock Neurotech, Salt Lake City, UT). All neural data streams were aligned to the behavioral data using a millisecond timestamp issued every 100ms by LiCoRICE[1] over a serial line aligned against the 30kHz timestamp of the Blackrock Neural Signal Processor. This approach corrected for any potential temporal slip arising from the two data streams with independent clocks. Offline, local field potential (LFP) signals were then extracted from the raw 30 kHz recordings by band-pass filtering with a fourth-order, zero-phase Butterworth filter from 1 – 250 Hz, notch-filtering at 60 Hz, and downsampling to 1 kHz.

Within our dataset, for each of the included trials each day, we could then extract peri-move 1 kHz LFP signals from M1 into a $n_{ch} \times n_{ms}$ array where $n_{ch} = 96$ channels and $n_{ms} =$501 ms. The LFP signal for each (channel, trial) pair was then converted to a periodogram using `scipy.signal.periodogram`[2] with a Hann window, "spectrum" scaling, and otherwise default parameters. Only the resultant coefficients for frequencies between 1 – 100 Hz were kept (of which there were 50). The median periodogram was then taken across all channels and trials for each day. Finally, from these daily periodograms, coefficients could then be partitioned across canonical frequency bands ($\delta$: 1 – 4 Hz,$\theta$: 4 – 8 Hz, $\alpha$: 8 – 12 Hz, $\beta$: 12 – 30 Hz, $\gamma$: 30 – 100 Hz) and summed to produce bandpower features.

For single-channel analyses, periodograms were computed by taking the median periodogram across trials for a given day. To remove noisy channels, single-channel periodograms with $\delta >$300 μV$^2$ were discarded. For periodograms from pretrial periods, these were computed identically but by considering the 500 ms before the radial reach cue was presented (501 ms total). This epoch of time contained the monkey returning the cursor to the center of the screen after a previous radial reach, and holding still at the center (Figure S4A). The same bandpower features were then computed for both single-channel and pretrial periodograms.

Similarly, for each (channel, trial) LFP signal on a given day, the continuous wavelet transform (CWT) implementation from PyWavelets[10] was used to estimate time-frequency dynamics. In particular, a complex Morlet wavelet[11] was used with previously reported parameters (center frequency: 1 Hz, bandwidth: 2 Hz).[12] This generated wavelet coefficients from 8 – 100 Hz in increments of 1 Hz (i.e. $n_f$=93 frequency bins) and in 10 ms temporal increments. To remove edge effects, the first and last three time points (i.e. 30 ms) were discarded from each time-frequency matrix, yielding $n_t$=45 time bins spanning 120 ms before to 320 ms after $T_{\text{rxn}}$. The median squared magnitude of each CWT coefficient was then taken across channels for each trial, and across trials for each day. This yielded a time-frequency tensor across $n_d$ days: $\mathcal{T} \in \mathbb{R}_+^{n_d \times n_f \times n_t}$ (Figure 4A).

### Analysis and modeling

#### Kinematic entropy

On each recording day, arm positions ($p_x$, $p_y$; mm) and velocities ($v_x$, $v_y$; cm s$^{-1}$) in the 350 ms after $T_{\text{rxn}}$ were collected. If there were 100 included trials for a given movement condition, then this gives 35,000 position/velocity estimates. For each condition, these estimates were then binned into 2D histograms, with 100 bins along both the $x/y$ dimensions spanning $\pm$110 mm/80 mm for Monkey U/H's position and $\pm$ 60 cm s$^{-1}$ for each monkey's velocity. This yielded a discrete position/velocity distribution for each *condition*. These were then averaged to yield a discrete position/velocity distribution for each *day*; these are essentially 10,000-dimensional (100$\times$100) vectors that sum to one, and examples are visualized in Figure S2A. This two-stage process of estimating condition-specific histograms before averaging ensures that histograms are not biased by differences in the number of reaches included per condition each day. Next, the entropy (in nats) of the position/velocity distributions was computed by Equation 1, yielding a position entropy ($H_p$) and velocity entropy ($H_v$) each day (Figure S2B).

$$H(p) = \sum_{i=1}^{10,000} p(i) \ln p(i) \tag{1}$$

#### Spectral parametrization

The "fitting oscillations and one-over-$f$" package (FOOOF) was used for spectral parametrization. For each animal, a model selection procedure was used to select key hyperparameters such as the maximum number of peaks; this is important, as permitting additional peaks can lead to over-fitting of noisy spectra. As pointed out in past work, the algorithm performs a multi-stage, nonlinear least-squares fit of the $\log_{10}$ empirical spectrum, and thus approximates a maximum likelihood solution under the assumption of additive, zero-mean Gaussian noise; as such, a Gaussian likelihood function can be used to compare model fits across different parametrizations.[13] This likelihood of a model $f$ with parameters $w$ for a spectrum $y$ with frequency bins $x$ is reproduced in Equation 2. The noise variance $\sigma^2$ was approximated as the model's residual variance.

$$-\ln p(y \mid x, w, \beta) = \frac{1}{2\sigma^2} \sum_{i=1}^{N} \{f(x_i, w) - y_i\}^2 + \frac{N}{2} \ln\left(2\pi\sigma^2\right) \tag{2}$$

The Akaike information criterion (AIC)[14] was used to compare models using either a "fixed" or "knee" aperiodic mode, as well as 1 – 10 maximum peaks. The AIC was computed for each parameter setting and for each daily spectrum; Figure S3A shows the median and IQR AIC across spectra. As shown in Figure S3A, the additional knee parameter did not improve the fit for Monkey U, and provided a marginal improvement for Monkey H (although with high variance across days). Fitting a knee for Monkey H also produced less consistent identification of periodic features and more variable $R^2$ (not shown), and thus a fixed mode was used for each animal. AIC tended to saturate at 3 peaks for Monkey U and 5 peaks for Monkey H, and these were selected as the maximum number of peaks. In addition, peak width limits were set to 2 – 12 Hz and a peak threshold of 1.5 standard deviations (SDs) was used; other parameters were left to their defaults. Using these hyperparameters, FOOOF represented each daily spectrum as an aperiodic exponent ($ap_{exp}$) and offset ($ap_{off}$), alongside the locations, heights, and widths of oscillatory peaks. The $ap_{exp}$ is a slope in logµV$^2$-logHz space and is dimensionless, while $ap_{off}$ captures the corresponding intercept (in logµV$^2$) and peak powers capture logµV$^2$ units of oscillations above the aperiodic component. Note that to minimize any impact of the 60 Hz notch filter, spectrum coefficients between 53 – 67 Hz were masked and linearly interpolated in log-log space for all model fits. The resultant models all exhibited high goodness-of-fit (median $R^2$ across days [IQR]; Monkey U: 0.9947 [0.9941, 0.9952]; Monkey H: 0.9977 [0.9960, 0.9987]) which was not affected by the lesions (Figure S3C-D).

To analyze estimated peaks, it was useful to identify oscillations that consistently reappeared over time in a similar frequency range. As such, the peak locations across all recording sessions were arranged in a vector for each monkey, and a 1D Gaussian mixture model (GMM) was fit with 3 (Monkey U) or 5 (Monkey H) clusters in scikit-learn.[15] The best model after 100 initializations was selected to partition peaks; peaks that were beyond 3 SDs of all components were considered outliers (“x” markings in Figure S3B). Using this procedure, two sensorimotor rhythms (SMRs) in the $\alpha$-$\beta$ range were consistently identified in each animal, denoted $SMR_{\downarrow}$ (days identified; Monkey U: 102/102; Monkey H: 58/70) and $SMR_{\uparrow}$ (days identified; Monkey U: 102/102; Monkey H: 69/70). These SMRs had peak frequencies $SMR^f_{\downarrow}$ (median [IQR]; Monkey U: 15.2 [14.9, 15.7]Hz; Monkey H: 21.2 [20.8, 21.7]Hz) and $SMR^f_{\uparrow}$ (Monkey U: 32.8 [32.1, 33.6]Hz; Monkey H: 34.8 [34.1, 36.9]Hz) that aligned with previously observed $\sim$20/30Hz SMRs in the macaque motor cortex[16] (Figure S3A). The widths (in Hz) of these peaks were also analyzed ($SMR^W_{\downarrow}$, $SMR^W_{\uparrow}$). Note that oscillations in the $\delta$, $\gamma$, and upper $\gamma$ ($\gamma_{\uparrow}$) ranges were also identified by the GMM (Figure S3A), but were not individually analyzed further due to either inconsistent identification or collisions with the 60 Hz notch filter from LFP preprocessing. The SMRs were not identified every day, perhaps due to fitting instabilities or the inability of the model to resolve adjacent peaks to consistent locations,[17] and thus a more robust summary of SMR-related activity was also computed on each recording day. To do so, the periodic spectrum in linear µV$^2$ units was formed from the estimated periodic parameters, and the total "bandpower" spanning 8 – 45 Hz was stored; this aggregate statistic was denoted $\sum SMR$. This summed range of periodic power is highlighted in gray in both Figure 2E and Figure S3A.

For single-trial and pretrial periodograms, spectral parametrization was also used to estimate aperiodic ($ap_{\text{off}}$, $ap_{\text{exp}}$), periodic ($\sum SMR$), and diagnostic ($R^2$) features with the same hyperparameters as above.

#### Movement-related beta activity

One representation of time-frequency dynamics, as presented in prior literature, focuses on movement-related $\beta$ dynamics such as the movement-related beta decrease (MRBD) and the post-movement beta rebound (PMBR).[18,19] From inspecting time frequency dynamics (e.g. left of Figure 4A), these temporal dynamics were most visible in the frequency range around $SMR^f_\uparrow$ in accordance with previous work showing that higher-frequency SMRs are more movement-responsive.[20] This is especially relevant to extracting MRBD/PMBR dynamics from the short, rapid movements contained in these 500 ms epochs. Accordingly, average wavelet power (arbitrary units) in 8 Hz bands surrounding median $SMR^f_\uparrow$ (Monkey U: 29 – 37 Hz; Monkey H: 31 – 39 Hz) was extracted, and denoted $\beta_{\text{tf}}$ (see Figure S8A). Baseline power was averaged from −120 – −100 ms (relative to $T_{\text{rxn}}$), yielding $\beta_{\text{ref}}$. Then, minimum baseline-normalized activity from −100 – 200 ms was MRBD while the maximum baseline-normalized activity from 200 ms onward was the PMBR. In these measurements, being less than or greater than 1 indicates reduction or amplification of power with respect to baseline. In these short behavioral windows, PMBR measurements did not necessarily exceed 1 (see Monkey U in Figure S8A-B), although MRBD was always below 1 as expected (Figure S8B).

#### Tensor decomposition

The canonical polyadic decomposition (CPD) represents a tensor as a sum of rank-one component tensors (Figure 4A).[21–23] In our design, a rank $R$ CPD yielded three factor matrices (each with

$R$ columns) across days frequency and time: $\mathcal{M} = [\![\boldsymbol{\tau}^d, \boldsymbol{\tau}^f, \boldsymbol{\tau}^t]\!]$. These factors were able to approximate the time-frequency tensor described previously as: $\mathcal{T} \approx \sum_{r=1}^{R} \tau_r^d \circ \tau_r^f \circ \tau_r^t$, where $\circ$ is a vector outer product between the $r$-th component factors.[23] A critical property of the CPD is that under a fairly weak condition, where the factor matrices $\boldsymbol{\tau}^d, \boldsymbol{\tau}^f, \boldsymbol{\tau}^t$ are full rank, it has a unique global optimum up to permutations and scaling.[24,25] This contrasts widely-used techniques such as principal component analysis (PCA) where it is necessary to impose additional constraints on factors, such as orthogonality, to find a unique low-rank approximation to data. This means that, aside from the unlikely situation where the underlying data is generated by orthogonal factors, PCA loadings cannot be confidently interpreted.[26] As such, interpretation of literal CPD factors is more principled than in the matrix decomposition case. All that's left is to address remaining invariance classes; to resolve scaling invariance, it is common to absorb the magnitude of all component factors for a given rank $r$ into a scaling factor $\lambda_r$,[23] expanding our model specification to $\mathcal{M} = [\![\boldsymbol{\lambda}; \boldsymbol{\tau}^d, \boldsymbol{\tau}^f, \boldsymbol{\tau}^t]\!]$. To resolve permutation invariance, factors were manually sorted within each animal to facilitate visual comparison.

However, a tradeoff is that the optimization problem is nonconvex and doesn't have a closed form solution (like in PCA); an alternating least squares approach is common, although this can get stuck in local minima and fail to recover the global optimum.[23,27] Moreover, the optimal solution can completely change depending on the number of components, meaning that each rank must be considered as its own optimization.[28] But, it has been shown both theoretically[29,30] and empirically[26] that non-negative decompositions produce more stable solutions and are more likely to uncover the optimal decomposition across multiple initializations. This motivated our construction of a non-negative tensor of CWT coefficient magnitudes, and thus we employed a multiplicative, non-negative optimization algorithm implemented in TensorLy.[31]

We then performed model selection within each animal to determine the model rank $R$, using a mixture of reconstruction score, cross-validation, and model similarity heuristics that have shown success in spiking neural data.[26] In particular, we swept model rank $R$ from 1 to 10, and randomly initialized and optimized 1000 models at each rank. Each optimized model yielded a tensor reconstruction $\widehat{\mathcal{T}}$ that was evaluated by a normalized reconstruction score, taking on values between 0 and 1 (Equation 3):

$$\text{score} = 1 - \frac{||\mathcal{T} - \widehat{\mathcal{T}}||^2_{\mathcal{F}}}{||\mathcal{T}||^2_{\mathcal{F}}} \tag{3}$$

These reconstruction scores are visualized in Figure 4B, and have similar heuristic value to explained variance curves in PCA. The model with the best score at each rank was a candidate model, and we additionally examined how similar each set of models was to the candidate model optimized at each rank. We computed its similarity to all 999 non-candidate models (shown in Figure 4B) as well as the mean similarity of the 99 next-best models ("+" markings in Figure 4B). This allowed us to assess whether most (or at least the best) initializations converged to a similar solution; if so, then this solution was likely close to the global optimum. This is a critical metric for determining whether one can reap the interpretation benefits of the CPD. The actual similarity metric was a multiplicative cosine similarity between factors and has been used in past work.[26,27] In particular, when comparing two models $\mathcal{M}$ and $\tilde{\mathcal{M}}$, we measured the maximum angular similarity across all possible factor permutations ($\Omega$) as shown in Equation 4. A threshold of 0.8 has been found to yield similar models in past work.[26]

$$\max_{\omega \in \Omega} \frac{1}{R} \sum_{r=1}^{R} \left[ \left( 1 - \frac{\left| \lambda_r - \tilde{\lambda}_{\omega(r)} \right|}{\max(\lambda_r, \tilde{\lambda}_{\omega(r)})} \right) \right] \left( \boldsymbol{\tau}^{d\top}_r \tilde{\boldsymbol{\tau}}^d_{\omega(r)} \cdot \boldsymbol{\tau}^{f\top}_r \tilde{\boldsymbol{\tau}}^f_{\omega(r)} \cdot \boldsymbol{\tau}^{t\top}_r \tilde{\boldsymbol{\tau}}^t_{\omega(r)} \right) \tag{4}$$

We also paired these diagnostics with 2-fold cross-validated reconstruction error, where models were fit with a random 50% of tensor entries masked.[26,31] Within each fold, the best model (on the masked data) was found across 10 random initializations, and then reconstruction score was then computed on the masked elements. The cross-validation was repeated 25 times; across 2 folds, this yielded 50 cross-validated reconstruction scores at each rank.

Based on all model selection information, such as saturation in reconstruction scores (original and cross-validated) and drop-offs in model similarity (particularly among the 100 best models), a rank ($R$) of 4 was selected for Monkey U and a rank of 3 was selected for Monkey H (Figure 4B).

For instance, Monkey U's mean model similarity dropped from 0.78 (0.87 across top 100) at $R = 4$ to 0.72 (0.74) at $R = 5$. Similarly, Monkey H's mean model similarity dropped from 0.83 (0.90 across top 100) at $R = 3$ to 0.76 (0.79) at $R = 4$.

**State space modeling**

Finally, a linear Gaussian state space model (LG-SSM) estimated the temporal extent of lesion effects.[32] In the model, observations of the features on a given day $t$ formed the vector $y_t$ for $t \in [1, T]$, and were modeled as arising from a latent state $x_t$ with linear dynamics. Lesions were a vector input $u_t$, where the $i$th value $u_{ti}$ is 1 when it is $i$ days after a lesion and 0 otherwise. We considered 5 day lesion effects. The model parameters ($\Theta = \{\boldsymbol{\mu}_1, \Sigma_1, A, Q, C, D, R\}$) included the initial state mean/covariance ($\boldsymbol{\mu}_1/\Sigma_1$), the linear dynamics/covariance ($A/Q$), the linear mapping from the state variables to the observation mean ($C$), the mapping from inputs to observations ($D$), and the observation covariance ($R$). The input matrix $D$ was of principal importance, where the value $d_{ij}$ captured the impact of a lesion after $j$ days on feature $i$. When considering the 5 inputs for a given features, these are referred to elsewhere in the text as $d_1, d_2, \ldots, d_5$. This model is specified in Equation 5 and is shown graphically in Figure 4A.

$$
\begin{aligned}
x_1 &\sim \mathcal{N}(\boldsymbol{\mu}_1, \Sigma_1) \\
x_t &\sim \mathcal{N}(Ax_{t-1}, Q) \quad z \in [2, \ldots, T] \\
y_t &\sim \mathcal{N}(Cx_t + Du_t, R)
\end{aligned}
\tag{5}
$$

All model parameters ($\Theta$) were estimated using gradient descent for 500 iterations in Dynamax[32] using the Adam optimizer with a learning rate of 0.01 and otherwise default parameters from Optax.[33] Three models were fit to Behavior ($T_{\text{rxn}}, v_{\text{reach}}, \theta_{\text{err}}$), Periodogram ($\delta, \theta, \alpha, \beta, \gamma, ap_{\text{exp}}, ap_{\text{off}}, \sum SMR$), and Tensor ($\tau_1^d, \tau_2^d, \ldots, \tau_R^d$) datasets. Datasets were $z$-scored before model fitting to facilitate comparison of downstream statistics. Moreover, each model's latent dimension was selected as the number of principal components needed to explain 90 % of the $z$-scored dataset's variance. Each dataset spanned $T_{\text{total}}$ days, but the corresponding features may have only been completely observed on $T \leq T_{\text{total}}$ days due to missing recording days. These $T$ days were treated as contiguous samples to facilitate modeling. Finally, parameters were estimated across 20 random initializations to ensure identifiability (particularly of $D$) across repeated fits; in general, entries of $D$ had small SD across initializations (seeTable S6, Table S7, and Table S8). Each of these initializations also provided an estimate of the marginal log likelihood of the dataset (denoted $SSM_{LL}$); this was also largely consistent across initializations (Table S5). However, to account for parameter estimation uncertainty, the mean $SSM_{LL}$ and the mean $D$ matrices across initializations were considered the core test statistic (rather than using values from a single random seed). These statistics were then tested for significance using permutation tests (see below).

#### Statistics

##### Permutation tests

A statistical throughline for all analyses is the use of nonparametric permutation tests to support claims.[34–36] This approach is a data-efficient way to discern whether lesion-induced effects could arise from the empirical fluctuations of recorded time series, or must be explained as a genuine lesion response. For each animal, *n* lesions were sparsely interleaved across a longitudinal dataset with the discretion of investigators. The null hypothesis motivating the permutation tests below is that the observed data would be equally likely if the lesions had occurred on any *n* random days; in other words, observed effects were just a result of the selected temporal position of lesion days, and not the lesions themselves. This supposes that lesion/non-lesion labels are exchangeable. Such an approach is conceptually similar to randomization testing, although this would require that the lesion days were formally randomized *a priori*, which was not the case in this dataset. This approach is scoped to make within-monkey inferences (e.g. “X occurred in response to lesions *in Monkey Y*”), enabling this study to use an ethical sample size of 2 monkeys as is standard in primate motor systems neuroscience. Importantly, within-subject permutation testing is distinguished from global inferences about a population; this would require taking a truly random sample of said population (e.g. measuring post-lesion perturbations from a random sample of rhesus macaques). As such, the statistics reported are not accompanied by confidence intervals (although percentile intervals of permuted null statistics are reported where helpful). However, it's worth noting that when attached to a broader context, local inference made from available data[37] can support a more general result, or can at least motivate the expense of mounting larger, data-intensive experiments. This latter sense of value is particularly relevant to this study which hopes to motivate the use of electrolytic lesions in additional primates to explore how brain networks and motor behavior respond to neuron loss of different scales.

**Lesion sensitivity**

To estimate whether lesions had a significant next day effect on a given feature within an animal, longitudinal time series were converted to $z$-scored first differences. The $n_L$ feature differences which straddled electrolytic lesions were extracted and averaged into a test statistic $\langle \Delta \rangle_L$. Because periodic features (e.g. $SMR_{\downarrow}^{P}$ and $SMR_{\uparrow}^{P}$) were not necessarily recovered from each daily spectrum, $n_L$ reflects the number of lesions for which the feature was present on both the day before and after the lesion. These counts are reported in Table S2. The null hypothesis was that $\langle \Delta \rangle_L$ reflected only the selected temporal positions of these $n_L$ differences rather than a consequence of the lesions themselves.[35] To test this, a null distribution was constructed by permuting the positions of these $n_L$ lesions and and recomputing $\langle \Delta \rangle_L$ 10,000 times. A two-sided $p$-value was then estimated using the percentile method. This statistic was computed for behavioral, spectral parametrization, bandpower, time-frequency, and tensor features, as well as for all individual CWT coefficients. The grouping of these tests into families for multiple comparisons is shown in the Multiple Comparisons section; complete results are collected in Table S2.

This statistic was also computed on the spectral features computed from pretrial periodograms.

#### State space modeling

As described above, gradient descent estimated the maximum likelihood parameters $\Theta$ of an LG-SSM model. The matrix $D$ mediated the 5 day lesion input for a given dataset, and was thus a target for statistical inference. To account for uncertainty in parameter estimation, the statistics of interest were the averaged matrix entries $d_{ij}$ across 20 random initializations. The null hypothesis, similar to that used for $\langle\Delta\rangle_L$, was that the value of $d_{ij}$ was just a consequence of the temporal positions of the lesions within the dataset. Thus, a null distribution was constructed by permuting the position of lesion days, reconstructing the inputs $u_t$, and re-fitting the model using gradient descent. Each point in the null distribution was computed by randomly initializing and averaging 5 $d_{ij}$ estimates (this was reduced from 20 due to computational burden). This was repeated 1000 times overall, and then a two-sided $p$-value was estimated using the percentile method (Table S6, Table S7, and Table S8). An analogous approach was used to test whether overall model fit was improved by including the structured lesion input $u_t$. For each dataset, the mean $SSM_{LL}$ across 20 initializations was the test statistic. If the lesions generated relevant input signals, then the model trained using the structured inputs $u_t$ should outperform a model where the inputs were generated using permuted lesion times. To test this, a null distribution was generated via the permutation procedure described above. A one-sided $p$-value was computed using the percentile method to test whether the observed mean $SSM_{LL}$ was larger than expected under the null (see results in Table S5).

#### Behavioral correlation

Longitudinal relationships between neural and behavioral features were investigated via Spearman correlation. To ensure that detected relationships capture shared day-to-day fluctuations rather than shared longitudinal trend, the time series were differenced to reduce non-stationarity. Computed correlation values ($\rho$) and corresponding $p$-values are included in Table S9 and Table S10.

#### Recurrent lesion effects

For a given feature, the longitudinal time series was converted to $z$-scored first differences, and the magnitude of differences straddling lesions (denoted $|\Delta|$) were collected. Of the 22 behavioral/spectral features (Multiple Comparisons), only the features that were identified around all lesions were further analyzed. This was all 22 features for Monkey U, but 19 features for Monkey H ($SMR^{P}_{\downarrow}$, $SMR^{W}_{\downarrow}$, $SMR^{f}_{\downarrow}$ were not identified around Lesion 4). This let lesions be treated as repeated measures of effects on a matched set of features. Thus, the 22/19 $|\Delta|$ estimates were collected for each lesion, and the Friedman test statistic ($\chi^2$) alongside an exact, permutation-based $p$-value was computed using `scipy.stats.friedmanchisquare`. This is reported for each animal alongside Kendall's W as an effect size. If the Friedman test indicated a significant effect, pairwise post-hoc comparisons between lesions was conducted with the Wilcoxon signed-rank test (two-sided). The test statistic ($T$) alongside an exact, permutation-based $p$-value was computed using `scipy.stats.wilcoxon`. These are presented alongside the proportion of features that increased/decreased in magnitude between selected lesions. Based on these statistical results, Page's trend test[38] was used to test for a decreasing ordering of repeated measures with successive lesions; the test statistic ($L$) and an exact, permutation-based $p$-value was computed with `scipy.stats.page_trend_test`. In addition, Spearman correlation (two-sided) was computed between lesion number and the $\Delta$ features, producing $\rho$ and a corresponding $p$-value with `scipy.stats.spearmanr`.

#### Spatial lesion effects

The five bandpower and four spectral parametrization ($ap_{\text{off}}$, $ap_{\text{exp}}$, $\sum SMR$, $R^2$) features were computed on single-channel periodograms. These time series were converted to $z$-scored first differences, and the differences straddling lesions ($\Delta$) were collected. For a given feature, this provides up to 96 $\Delta$ features per lesion. To test whether $\Delta$ tended to increase/decrease based on spatial proximity to the lesion, Spearman correlation was computed between $\Delta$ estimates across all channels/lesions and Manhattan distance to the nearest lesion electrode. Electrodes had a pitch of 400 µm.

#### Multiple comparisons

The false discovery rate was used to control the family-wise error rate at 0.05 for each group of statistical tests.[39] For the $\langle \Delta \rangle_L$ statistic, the first family considered all static periodogram and behavioral features. These included bandpowers ($\delta, \theta, \alpha, \beta, \gamma$), spectral parametrization features ($ap_{\text{exp}}$, $ap_{\text{off}}$, $\sum SMR$, $SMR_{\downarrow}^{f}$, $SMR_{\uparrow}^{f}$, $SMR_{\downarrow}^{P}$, $SMR_{\uparrow}^{P}$, $SMR_{\downarrow}^{W}$, $SMR_{\uparrow}^{W}$), FOOOF diagnostics ($R^2$), behavioral features ($\text{rate}_{\text{suc}}$, $T_{\text{rxn}}$, $v_{\text{reach}}$, $\theta_{\text{err}}$,$H_p$, $H_v$), and behavioral diagnostics ($\text{rate}_{\text{acc}}$), totaling 22 tests. For pretrial periodograms, these $p$-values were corrected across 9 spectral features ($\delta, \theta, \alpha, \beta, \gamma$, $ap_{\text{exp}}$, $ap_{\text{off}}$, $\sum SMR$, $R^2$). Next, dynamic time-frequency features, introduced later in the text and analysis, were considered a separate family; these included $\beta_{\text{tf}}$ dynamics ($\beta_{\text{ref}}$, MRBD, PMBR) as well as tensor factors ($\tau_1^d, \tau_2^d, \ldots, \tau_R^d$), yielding 6 tests for Monkey H and 7 tests for Monkey U. CWT coefficients were also treated as their own family, yielding 4185 tests for each monkey. For LG-SSM statistics, each model was considered to produce its own family of tests. Thus, if a dataset had $k$ features, it produced a family of $k \times 5$ statistical tests for each entry $d_{ij}$ of the input matrix $D$. For instance, considering the model fit to the 8 periodogram features ($\delta, \theta, \alpha, \beta, \gamma$, $ap_{\text{exp}}$, $ap_{\text{off}}$, $\sum SMR$), these $p$-values were corrected across 40 tests. The singular marginal log likelihood statistic ($SSM_{LL}$) for each dataset was not corrected. For behavioral correlation statistics, each behavioral feature ($T_{\text{rxn}}$, $v_{\text{reach}}$, and $\theta_{\text{err}}$) produced its own family of tests. Moreover, two groups of covariates were corrected separately for each target: 14 static periodogram features ($\delta, \theta, \alpha, \beta, \gamma$, $ap_{\text{exp}}$, $ap_{\text{off}}$, $\sum SMR$, $SMR_{\downarrow}^{P}$, $SMR_{\downarrow}^{f}$, $SMR_{\downarrow}^{W}$, $SMR_{\uparrow}^{P}$, $SMR_{\uparrow}^{f}$, $SMR_{\uparrow}^{W}$) and time-frequency features (i.e. 3 $\beta_{\text{tf}}$ features plus $R$ tensor factors). For recurrent lesion effects, post-hoc pairwise Wilcoxon signed-rank tests were corrected using the false discovery rate. For $n$ lesions, this yields $\binom{n}{2}$ tests. Finally, spatial correlations were corrected across 9 spectral features ($\delta, \theta, \alpha, \beta, \gamma$, $ap_{\text{exp}}$, $ap_{\text{off}}$, $\sum SMR$, $R^2$).

#### Visualization

Plots of longitudinal trends (e.g. Figure 1E) include a smoothed visualization of the time series computed using Locally Weighted Scatterplot Smoothing as implemented in `statsmodels`[40] with a regression window of 5 days.

### Supplemental Figures

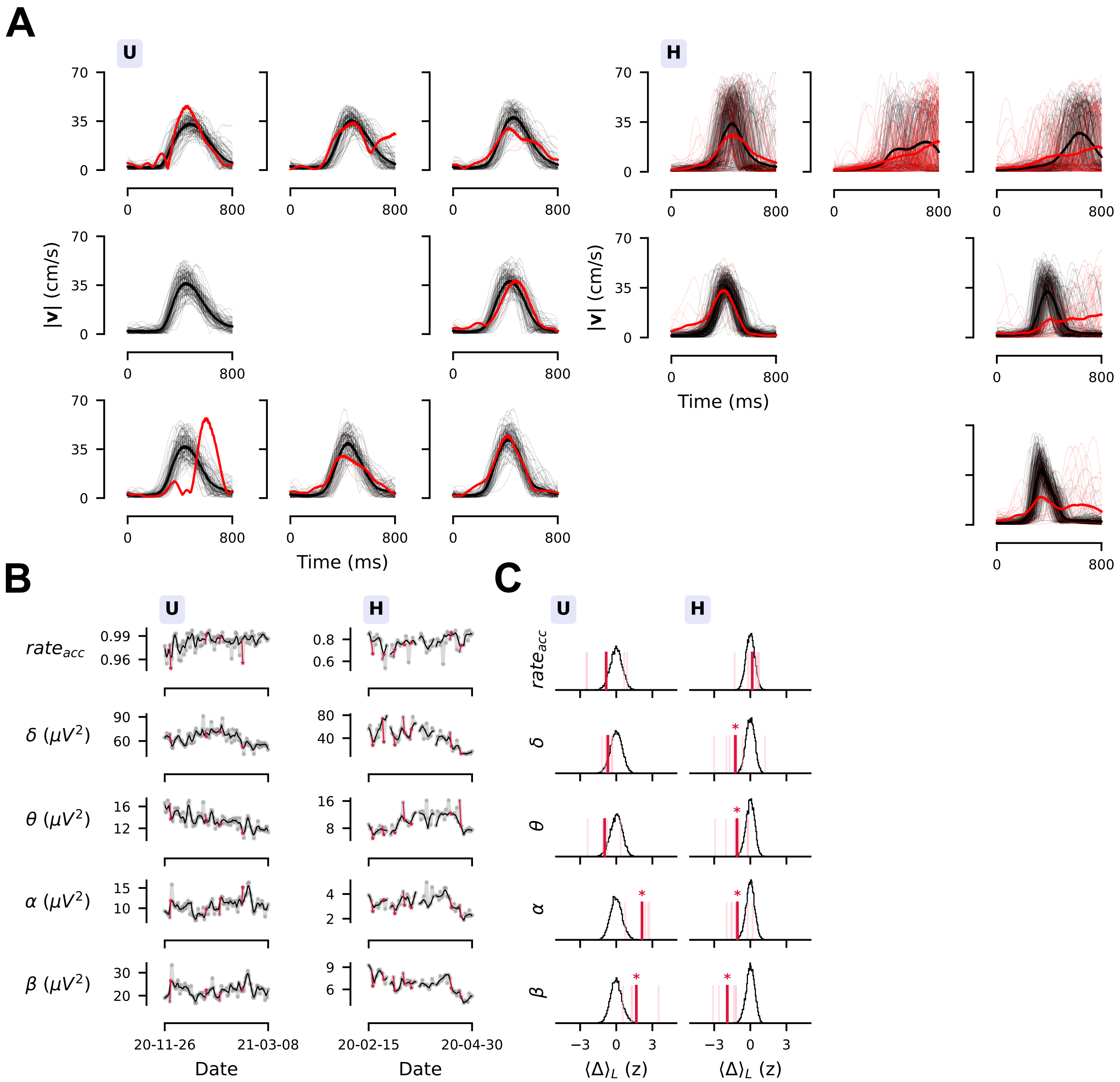


**Figure S1: Trial inclusion examples and additional features. (A)** Behavioral filters were applied to individual trials in order to only include stereotyped reaches (see Detailed Methods). Examples of excluded (red) and included (black) speed traces (thin traces are single trials, thick lines are means) relative to target presentation are shown for 2020-12-01 in Monkey U and 2020-04-18 in Monkey H. Note that reaches to the top and top right in Monkey H had delayed and highly variable reaction times that would often be rejected by behavioral filters; only 49.4 % of reaches to these targets were stereotyped while 88.0 % of reaches to other targets met the requirements. These two targets were excluded from analysis in order to isolate highly stereotyped behavior. In Monkey U, 98.0 % of reaches on this day were stereotyped. **(B-C)** Additional longitudinal trends (B) and lesion sensitivity statistics (C, $*p < 0.05$) for behavioral and bandpower features demonstrate that activation of behavioral filters was not affected by lesions ($\text{rate}_{\text{acc}}$). In addition, there was a divergent impact on $\alpha$ and $\beta$ bandpower which increased after lesions in Monkey U while decreasing in Monkey H. Additional effects included decreases in $\delta$ and $\theta$ for Monkey H. See full numerical statistics in Table S2.

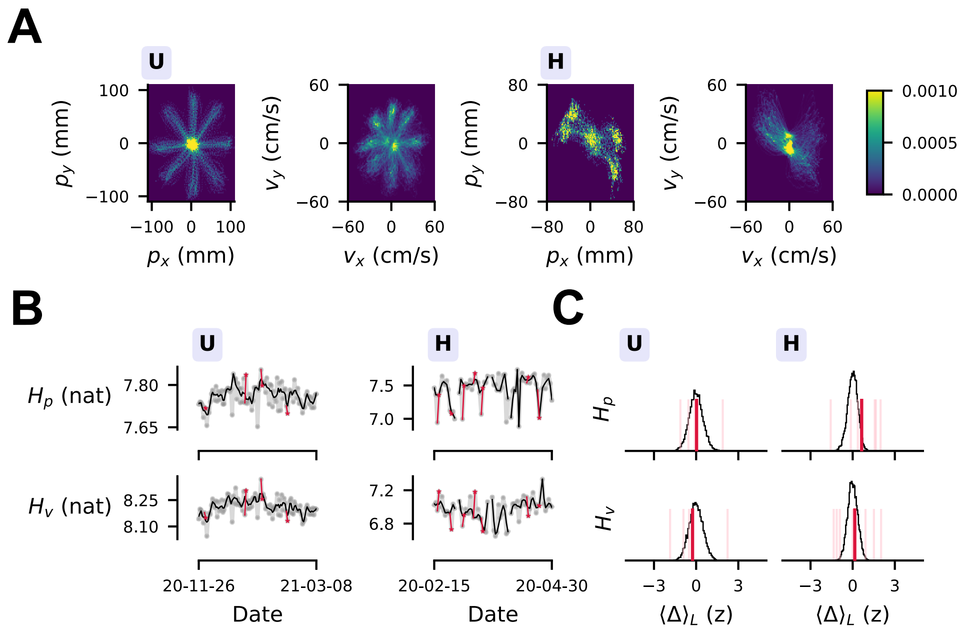


**Figure S2: Kinematic entropy analysis.** **(A)** Each day, 2D discrete distributions of position and velocity were computed (see Detailed Methods). The left two panels show Monkey U, within which the left/right panels show the position/velocity distribution for the date 2020-12-01. On the right are the position/velocity distribution for 2020-02-16 in Monkey H. These distributions were used to get daily estimates of position/velocity entropy ($H_p$ and $H_v$). **(B-C)** The longitudinal trend (B) and sensitivity (C) are shown for both $H_p$ and $H_v$. There was no effect of lesions on $H_p$ (Monkey U: $\langle\Delta\rangle_L = 0.03$, $p = 1.000$; Monkey H: $\langle\Delta\rangle_L = 0.64$, $p = 0.124$) or $H_v$ (Monkey U: $\langle\Delta\rangle_L = -0.26$, $p = 0.742$; Monkey H: $\langle\Delta\rangle_L = 0.15$, $p = 0.817$) (*$p < 0.05$), supporting the conclusion that neuron loss was covert. See full numerical statistics in Table S2.

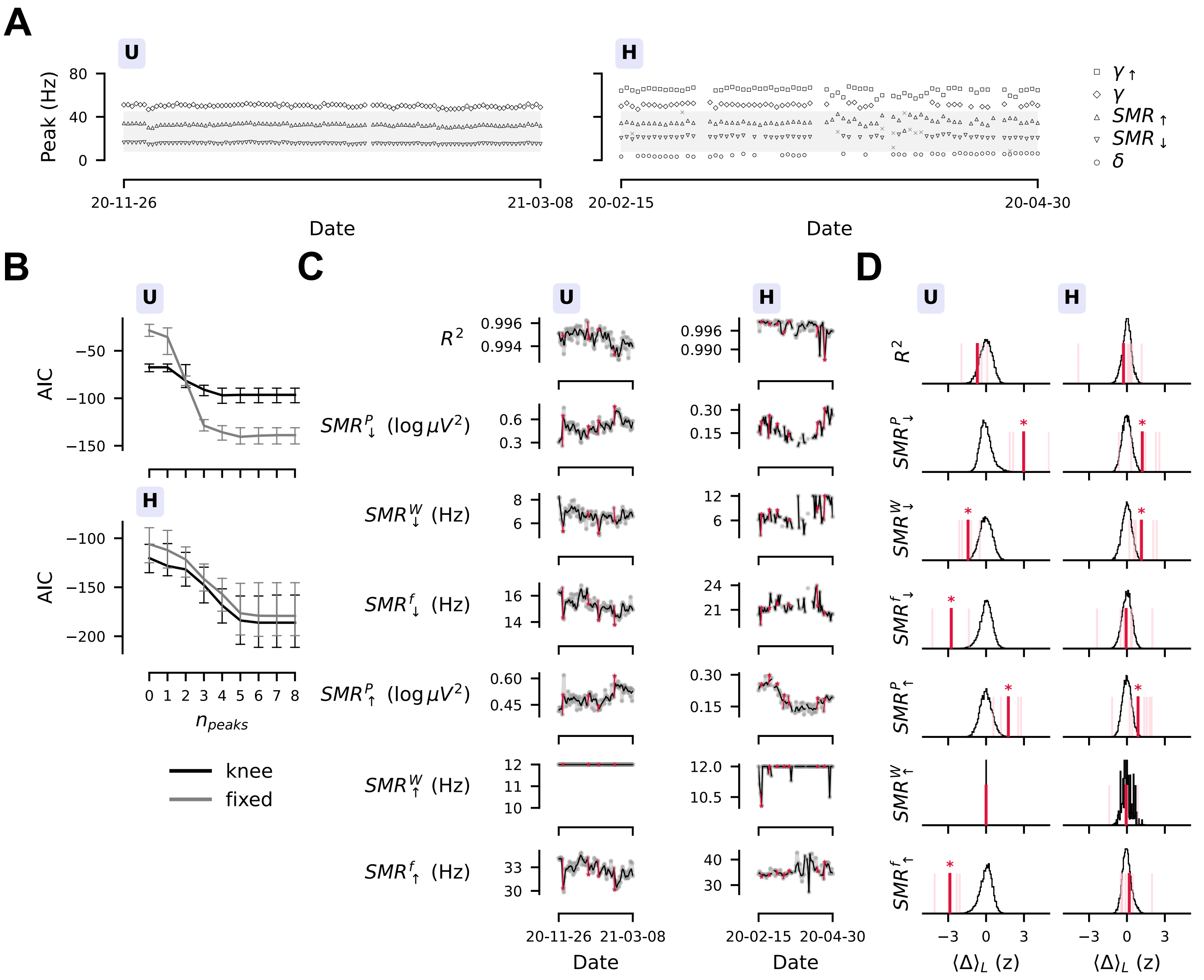


**Figure S3: Spectral parametrization model fitting**. **(A-B)** A model selection approach was used to determine the aperiodic mode and maximum number of peaks.[13] Median and IQR AIC across all daily spectra for each parameter setting is shown in B. Inclusion of an additional "knee" parameter was not well-motivated in either monkey, while AIC saturated at 3 peaks in Monkey U and 5 peaks in Monkey H. Using 3 (Monkey U) and 5 (Monkey H) maximum peaks, the detected peak locations are shown in A. Peaks had consistent structure over time; this was extracted using a GMM to group the oscillatory peaks into 3 (Monkey U) or 5 (Monkey H) bands; only $SMR_{\downarrow}$ and $SMR_{\uparrow}$ were further analyzed due to consistent identification, alignment with known SMRs in the macaque motor cortex,[16] and lack of collision with 60 Hz notch filtering. In addition, an aggregate periodic bandpower in $\mu V^2$ was computed from 8 – 45 Hz, denoted by the gray shading and denoted $\sum SMR$ in the main text. **(C-D)** Additional longitudinal trends (C) and lesion sensitivity statistics (D) found that goodness-of-fit ($R^2$) was not affected by lesions (*$p < 0.05$). In addition, SMR powers ($SMR^P_{\downarrow}$ and $SMR^P_{\uparrow}$) were both increased after lesions. $SMR^W_{\downarrow}$ increased in Monkey H but decreased in Monkey U, while $SMR^W_{\uparrow}$ was unaffected. In addition, center frequencies ($SMR^f_{\downarrow}$ and $SMR^f_{\uparrow}$) were reduced after lesions in Monkey U. See full numerical statistics in Table S2.

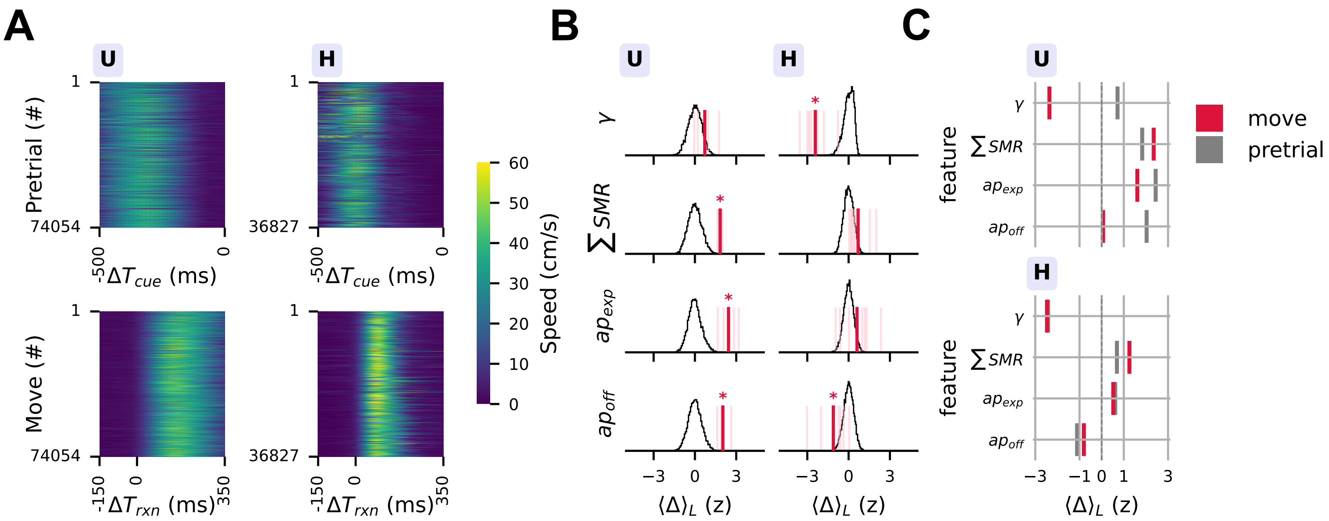


**Figure S4: Pretrial lesion effects. (A)** Speed traces for all included trials during the Pretrial phase (top) and the Move phase (bottom). The experimental flow did not include a movement-free inter-trial period, but the 500 ms before the reach stimulus was analyzed to probe the movement-specificity of lesion effects. Heatmaps are colored according to calculated arm speed in units of cm s$^{-1}$. **(B)** The main effects of $\gamma$ reduction and $\sum SMR$ amplification were not replicated in the pretrial period for both monkeys. In Monkey U, an elevated pretrial $ap_{\text{off}}$ prevented the movement-related $\gamma$ reduction shown in Figure 1E-F. In both monkeys, $\sum SMR$ still tended to increase (Monkey U: $\langle \Delta \rangle_L = 1.83$, $p < 1 \times 10^{-3}$; Monkey H: $\langle \Delta \rangle_L = 0.69$, $p = 0.060$) but did not reach significance in Monkey H. **(C)** The pretrial and movement-related lesion sensitivity effect size ($\langle \Delta \rangle_L$) showed that $\sum SMR$ effects were reduced in both monkeys when considering the pretrial period (pretrial vs move; Monkey U: $\langle \Delta \rangle_L = 1.83$ vs $\langle \Delta \rangle_L = 2.35$; Monkey H: $\langle \Delta \rangle_L = 0.69$ vs $\langle \Delta \rangle_L = 1.26$). Aperiodic effects were consistent in Monkey H but were more variable in Monkey U. See pretrial lesion sensitivity statistics in Table S3.

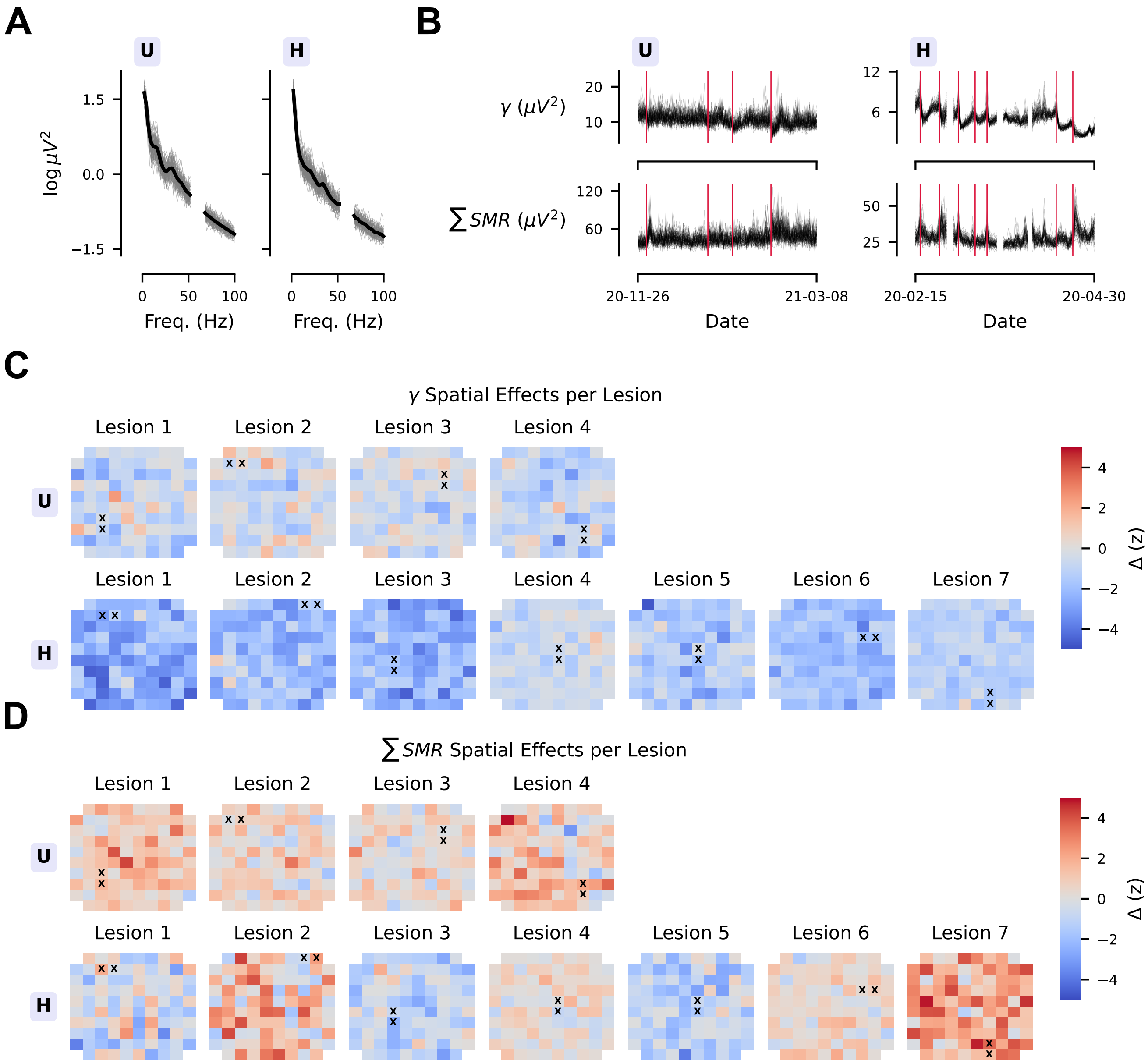


**Figure S5: Spatial distribution of lesion impact. (A)** Single-channel periodograms (thin grey traces) were noisier than the aggregated daily periodograms (thick black trace) used in all other analyses. Shown here are the periodogram estimates from near the start of the dataset: 2020-12-01 for Monkey U and 2020-02-16 for Monkey H. The 60 Hz notch filter is obscured from 53 – 67 Hz. **(B)** Lesion impact on single-channel $\gamma$ and $\sum SMR$ features was visible. The vertical crimson lines indicate lesion days, while the black traces show raw estimates of $\gamma$ and $\sum SMR$ features from single-channel periodograms. **(C)** For each analyzed lesion, the spatial distribution of $\gamma$ impact did not show clear structure. Heatmaps are colored according to $\Delta$, the $z$-scored difference in $\gamma$ from a pre-lesion session to another recording session ≈24 hours later (Figure 1C). $\gamma$ reduction was not concentrated around lesion electrodes, annotated with a black "x". The spacing between Utah array electrodes was 400 µm. **(D)** The same as (C) but for the spatial distribution of $\sum SMR$ impact; once again, there was no clear spatial structure to $\sum SMR$ impact. Results of Spearman correlation tests between $\Delta$ and Manhattan distance to nearest lesion electrode are visualized and reported in Figure S6 and Table S4.

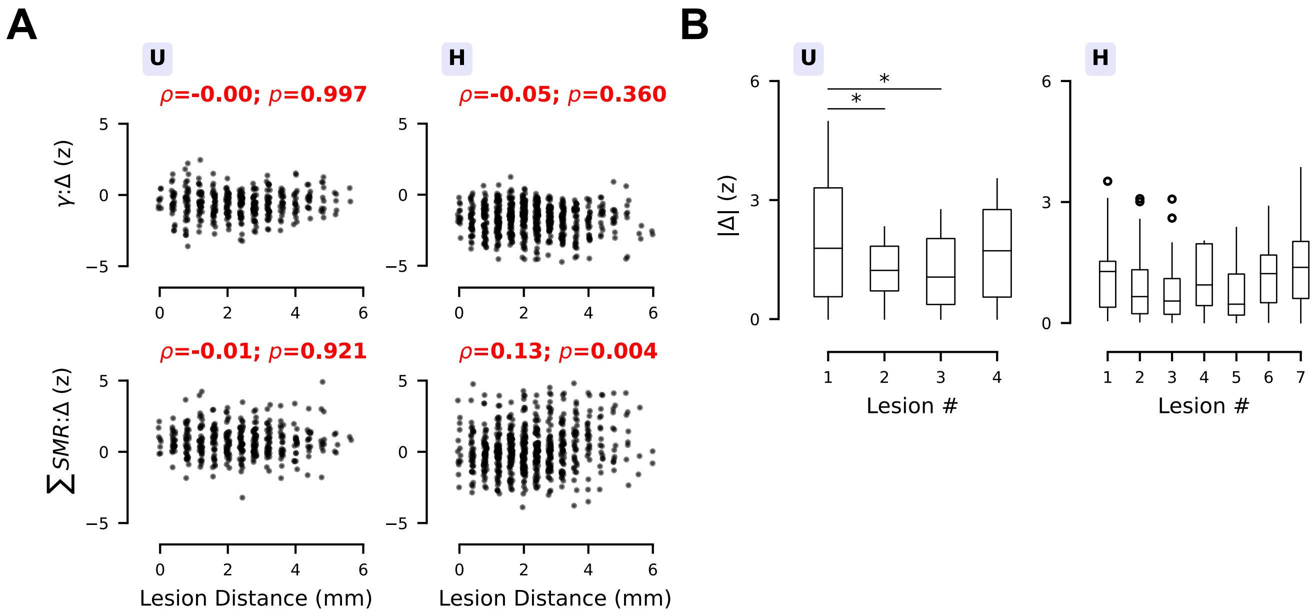


**Figure S6: Statistical outcomes of spatial and recurrent lesion analyses.** **(A)** Scatter plots of channel-wise $z$-scored first differences in $\gamma$ and $\sum SMR$ against Manhattan distance to nearest lesion electrode. For visibility, points are jittered by sampling from a uniform distribution on [−0.05, 0.05] mm. The impact on $\sum SMR$ in Monkey H had a a small but significant positive correlation ($\rho = 0.13$, $p = 0.004$, $N = 672$); no other correlation was significant. In correspondence with the diffuse spatial impact visualized in Figure S5C-D, all effect sizes were small ($\rho < 0.2$). See all spatial correlation statistics in Table S4. **(B)** Boxplots of effect size magnitude ($|\Delta|$) for features that were measured around each lesion (Monkey U: 22 features; Monkey H: 19 features). Boxplots show the median and IQR of these effects, while whiskers extend an additional 1.5 × IQR. In Monkey U, the Friedman test revealed a difference in effect size ($\chi^2(3) = 9.42$, $W = 0.14$, $p = 0.024$). Post-hoc pairwise Wilcoxon signed-rank tests found that Lesion 1 produced larger median effect sizes than Lesions 2/3 (1 vs 2: $T = 34$, 72.7 % decreased, $p = 0.024$; 1 vs 3: $T = 32$, 68.2 % decreased, $p = 0.024$). But, there was no monotonic decrease in effect sizes as tested with Page's trend test (Monkey U: $L = 561.0$, $p = 0.220$; Monkey H: $L = 2096.0$, $p = 0.742$) nor was there significant Spearman correlation between effect size and lesion position (Monkey U: $\rho = -0.06$, $p = 0.600$, $N = 88$; Monkey H: $\rho = 0.10$, $p = 0.260$, $N = 133$).

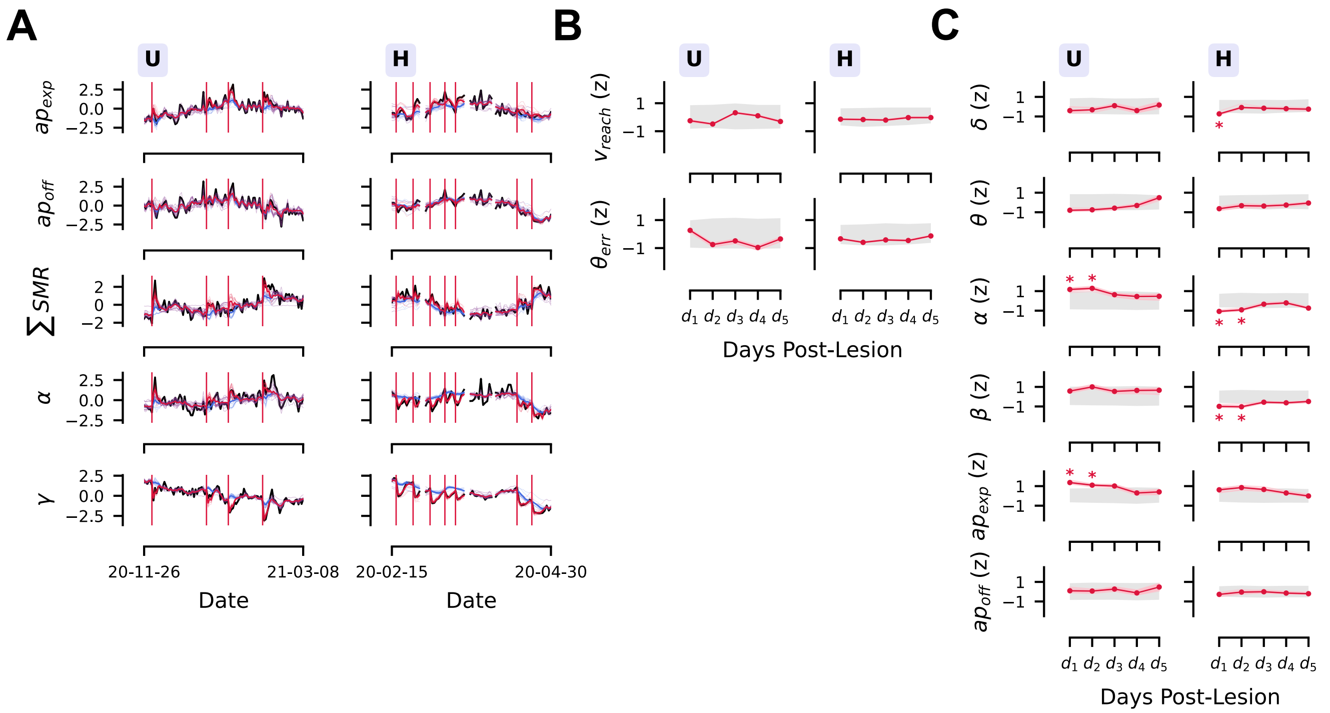


**Figure S7: Additional state space modeling. (A)** Smoothed reconstructions are visualized of selected periodogram features ($ap_{\text{exp}}$, $ap_{\text{off}}$, $\sum SMR$, $\alpha$, $\gamma$). Black lines indicate measured data, while thin lines indicates reconstructions from each of 20 model fits either including inputs (crimson) or without them (blue). Vertical crimson lines indicate lesions. Thick lines indicate mean reconstruction. **(B-C)** Individual lesion inputs are shown for additional features from the Behavior (B) and Periodogram (C) datasets; significance at $p < 0.05$ is indicated with a crimson asterisk. Null distributions were constructed using permutation testing (see Detailed Methods). Behavioral features shown here were unaffected by lesions according to the model, while additional effects on the periodogram features were uncovered. These included a 2 day increase to $ap_{\text{exp}}$ and a 2 day increase to $\alpha$ in Monkey U, alongside a 2 day decrease to both $\alpha$ and $\beta$ as well as a 1 day decrease to $\delta$ in Monkey H. These results underline that the spectrum is more sensitive to neuron loss than precise behavioral measurements. See all input parameters in Table S6 and Table S7.

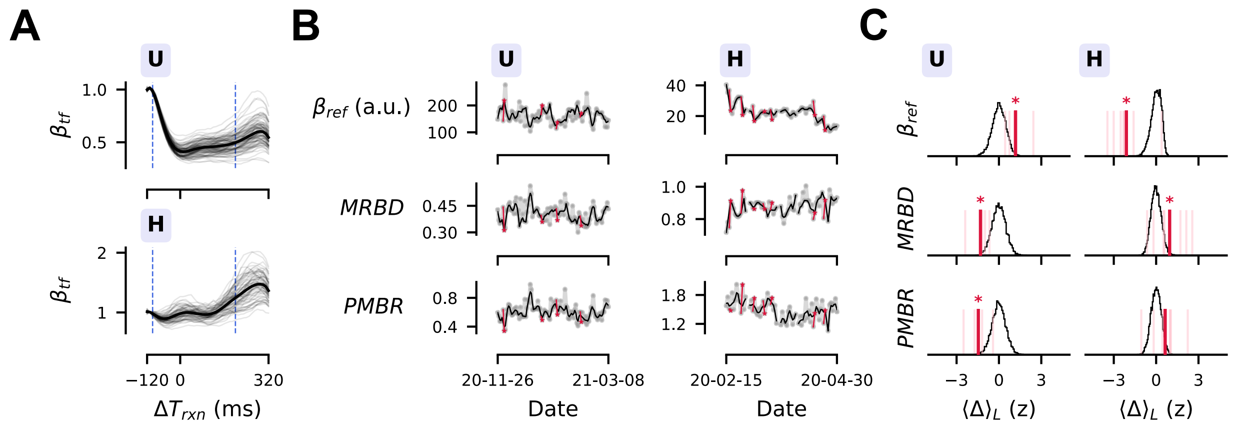


**Figure S8: Divergent lesion impact on movement-related $\beta_{\text{tf}}$ dynamics.** **(A)** While movement epochs were shortened, a partial MRBD/PMBR dynamic was visible when considering 8 Hz frequency bands centered on median $SMR_{\uparrow}^{f}$ in each monkey. **(B-C)** Longitudinal trends for baseline $\beta_{\text{tf}}$ power ($\beta_{\text{ref}}$), as well as normalized MRBD/PMBR are shown (B) alongside significance of the lesion sensitivity statistic $\langle\Delta\rangle_L$ (*$p < 0.05$). In accordance with divergent post-lesion impact on $\beta$ bandpower (see Figure S1B-C), $\beta_{\text{ref}}$ increased in Monkey U ($\langle\Delta\rangle_L = 1.18$, $p = 0.015$) but decreased in Monkey H ($\langle\Delta\rangle_L = -2.11$, $p < 1 \times 10^{-3}$) (C). With increased dynamic range, this increased the magnitude of (decreased the value of) MRBD in Monkey U ($\langle\Delta\rangle_L = -1.31$, $p = 0.005$), while the opposite occurred in Monkey H ($\langle\Delta\rangle_L = 0.958$, $p = 0.019$) (C). Additionally, PMBR decreased in magnitude (decreased in value) for Monkey U ($\langle\Delta\rangle_L = -1.46$, $p = 0.005$) (C).

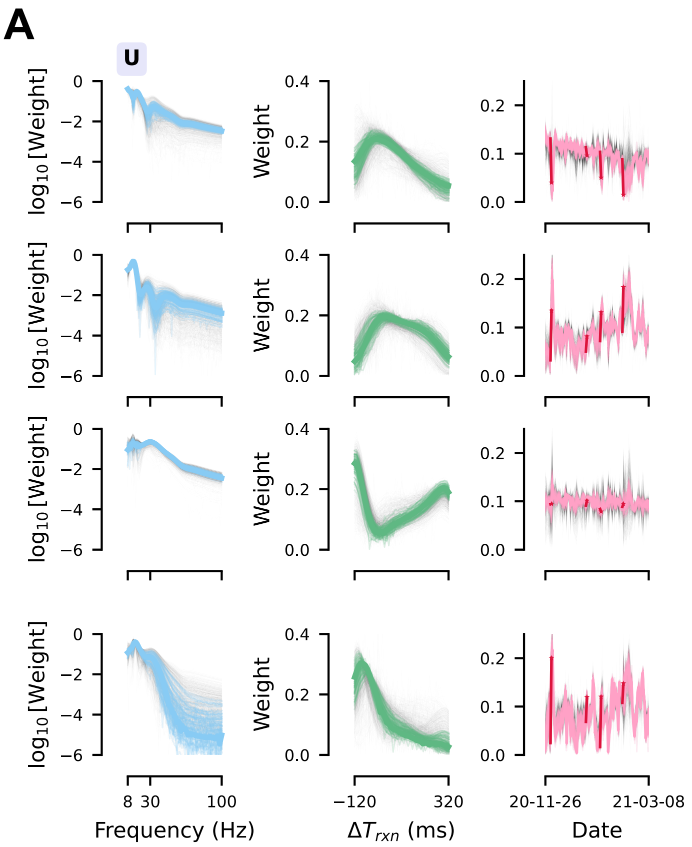


**Figure S9: Additional tensor decomposition visualizations. (A)** High similarity of estimated factors across repeated initializations can help ensure that the unique, globally optimal CPD has been identified.[24,26] Ranks of $R = 4$ and $R = 3$ were chosen for Monkeys U/H alike due to high model similarities across 1000 initializations relative to the highest scoring model, especially for the next 99 next-best models (see Figure 4B). Identified factors across all initializations are shown to indicate qualitative model similarity. The selected model is shown with a colored, thick, opaque trace. The 99 next-best models are shown in colored, thin traces. Remaining models are shown as gray, thin traces. When visualizing the selected date factors $\tau_r^d$, crimson markings indicate next-day impact of electrolytic lesions, just as in other longitudinal trend visualizations.

### Supplemental Tables

| | | $I_L$ ($\mu A$) | $T_L$ (s) | T ($^\circ F$) | pH | $pCO_2$ | $HCO_3$ | $TCO_2$ | BE | $pO_2$ | $SpO_2$ | Lac. | Arm |
|---|---|---|---|---|---|---|---|---|---|---|---|---|---|
| 19-12-03 | H | 150 | 45 | | | | | | | | | | 5 |
| 20-01-13 | H | 140 | 45 | | | | | | | | | | 5 |
| 20-01-20 | H | 130 | 50 | 103.0 | 7.455 | 27.8 | 19.6 | 20.0 | -4.0 | 85.0 | 97.0 | 2.14 | 5 |
| 20-02-17 | H | 160 | 45 | 99.9 | 7.468 | 37.8 | 27.4 | 29.0 | 4.0 | 69.0 | 95.0 | 0.89 | 4+ |
| 20-02-25 | H | 160 | 45 | 99.3 | 7.421 | 37.4 | 24.3 | 25.0 | 0.0 | 76.0 | 95.0 | 1.39 | 4+ |
| 20-03-04 | H | 170 | 45 | 99.2 | 7.500 | 29.7 | 23.1 | 24.0 | 0.0 | 103.0 | 99.0 | 1.41 | 5 |
| 20-03-11 | H | 160 | 45 | 99.1 | 7.443 | 34.3 | 23.5 | 24.0 | -1.0 | 77.0 | 96.0 | 0.93 | 5 |
| 20-03-16 | H | 160 | 45 | 98.7 | 7.423 | 41.7 | 27.2 | 28.0 | 3.0 | 64.0 | 92.0 | 0.99 | 5 |
| 20-04-14 | H | 150 | 45 | 98.6 | 7.425 | 39.6 | 26.0 | 27.0 | 2.0 | 67.0 | 94.0 | 1.03 | 4+ |
| 20-04-21 | H | 480 | 30 | 97.5 | 7.478 | 31.6 | 23.4 | 24.0 | 0.0 | 95.0 | 98.0 | 0.93 | 4+ |
| 20-12-01 | U | 170 | 30 | 100.6 | 7.535 | 30.7 | 26.0 | 27.0 | 3.0 | 124.0 | 99.0 | 2.16 | 5 |
| 21-01-05 | U | 170 | 30 | | 7.463 | 38.0 | 27.2 | 28.0 | 3.0 | 66.0 | 94.0 | 1.33 | 5 |
| 21-01-19 | U | 170 | 30 | | 7.492 | 36.8 | 28.2 | 29.0 | 5.0 | 88.0 | 97.0 | 1.20 | 5 |
| 21-02-10 | U | 160 | 30 | | 7.488 | 35.6 | 27.0 | 28.0 | 4.0 | 126.0 | 99.0 | 1.44 | 5 |

**Table S1: Lesion parameters and medical workup**. Lesion current ($I_L$) and duration ($T_L$) are shown. Blood gas measurements were made for all lesions except the first two in Monkey H (measured for all analyzed lesions). Partial pressures ($pCO_2$, $pO_2$) are in mmHg, concentrations ($HCO_3$, $TCO_2$, BE, Lac.) are in mmol $L^{-1}$, Arm is muscle strength grading of contralesional arm, while $SpO_2$ is in percentage. Blood tests and temperature, where available, were taken within two days of lesion occurrence, often on the same day. No significant abnormalities in blood gas or arm strength were observed for included lesions. Lesion parameters were chosen from a calibrated range of 150 – 170 µA and 30 – 45 s except for the third (130 µA, 50 s) and the exploratory final lesion (480 µA, 30 s) in Monkey H.

| | U | | | | H | | | |
|---|---|---|---|---|---|---|---|---|
| | $\langle\Delta\rangle_L$ (unit) | $\langle\Delta\rangle_L$ (z) | $p$ | $n_L$ | $\langle\Delta\rangle_L$ (unit) | $\langle\Delta\rangle_L$ (z) | $p$ | $n_L$ |
| $rate_{suc}$ | 0.000 | −0.012 | 1.0000 | 4 | −0.001 | −0.022 | 0.8279 | 7 |
| $rate_{acc}$ | −0.010 | −0.837 | 0.1581 | 4 | 0.016 | 0.158 | 0.8169 | 7 |
| $T_{rxn}$ $(ms)$ | 17.125 | 2.123 | 0.0006 | 4 | 4.071 | 0.282 | 0.6380 | 7 |
| $v_{reach}$ $(cm/s)$ | −1.001 | −0.650 | 0.2546 | 4 | 0.189 | 0.044 | 0.8626 | 7 |
| $\theta_{err}$ (deg) | 0.453 | 0.306 | 0.6833 | 4 | −4.402 | −0.547 | 0.2006 | 7 |
| $\delta$ $(\mu V^2)$ | −5.213 | −0.695 | 0.2480 | 4 | −16.816 | −1.239 | 0.0044 | 7 |
| $\theta$ $(\mu V^2)$ | −1.117 | −0.963 | 0.1082 | 4 | −2.790 | −1.107 | 0.0126 | 7 |
| $\alpha$ $(\mu V^2)$ | 3.401 | 2.138 | 0.0004 | 4 | −0.604 | −1.069 | 0.0126 | 7 |
| $\beta$ $(\mu V^2)$ | 4.323 | 1.669 | 0.0036 | 4 | −1.351 | −1.911 | 0.0011 | 7 |
| $\gamma$ $(\mu V^2)$ | −1.384 | −2.359 | 0.0004 | 4 | −1.661 | −2.469 | 0.0011 | 7 |
| $ap_{exp}$ | 0.051 | 1.612 | 0.0034 | 4 | 0.042 | 0.523 | 0.2484 | 7 |
| $ap_{off}$ $(\log \mu V^2)$ | 0.003 | 0.063 | 1.0000 | 4 | −0.097 | −0.806 | 0.0788 | 7 |
| $SMR^P_\downarrow$ $(\log \mu V^2)$ | 0.234 | 2.977 | 0.0004 | 4 | 0.069 | 1.212 | 0.0066 | 6 |
| $SMR^P_\uparrow$ $(\log \mu V^2)$ | 0.078 | 1.764 | 0.0022 | 4 | 0.028 | 0.878 | 0.0323 | 7 |
| $SMR^f_\downarrow$ (Hz) | −1.482 | −2.759 | 0.0004 | 4 | −0.070 | −0.065 | 0.8626 | 6 |
| $SMR^f_\uparrow$ (Hz) | −2.665 | −2.863 | 0.0004 | 4 | 0.651 | 0.183 | 0.7488 | 7 |
| $SMR^W_\downarrow$ (Hz) | −0.812 | −1.428 | 0.0035 | 4 | 2.696 | 1.150 | 0.0066 | 6 |
| $SMR^W_\uparrow$ (Hz) | 0.000 | 0.000 | 1.0000 | 4 | −0.035 | −0.066 | 0.8626 | 7 |
| $\sum SMR$ $(\mu V^2)$ | 8.682 | 2.352 | 0.0004 | 4 | 2.404 | 1.255 | 0.0044 | 7 |
| $R^2$ | −0.001 | −0.693 | 0.2487 | 4 | −0.001 | −0.275 | 0.6380 | 7 |
| $H_p$ (nat) | 0.003 | 0.031 | 1.0000 | 4 | 0.155 | 0.640 | 0.1244 | 7 |
| $H_v$ (nat) | −0.015 | −0.256 | 0.7419 | 4 | 0.013 | 0.150 | 0.8169 | 7 |
| $\tau^d_1$ (a.u.) | −0.059 | −2.713 | 0.0002 | 4 | −0.042 | −1.470 | 0.0003 | 7 |
| $\tau^d_2$ (a.u.) | 0.071 | 2.615 | 0.0002 | 4 | −0.003 | −0.150 | 0.6682 | 7 |
| $\tau^d_3$ (a.u.) | 0.003 | 0.290 | 0.5224 | 4 | −0.025 | −1.490 | 0.0004 | 7 |
| $\tau^d_4$ (a.u.) | 0.093 | 2.011 | 0.0002 | 4 | | | | |
| $\beta_{ref}$ (a.u.) | 36.382 | 1.178 | 0.0152 | 4 | −8.341 | −2.107 | 0.0003 | 7 |
| $MRBD$ | −0.071 | −1.308 | 0.0053 | 4 | 0.066 | 0.958 | 0.0192 | 7 |
| $PMBR$ | −0.171 | −1.460 | 0.0049 | 4 | 0.178 | 0.641 | 0.0864 | 7 |

**Table S2: Lesion sensitivity statistics**. $\langle\Delta\rangle_L$ is shown original units and in $z$-score (as reported in the text) alongside $p$-values from permutation testing. $n_L$ refers to the number of lesions where this feature was observed before/after the lesion (oscillatory peaks were not always identified on each recording day, although aggregate SMR power $\sum SMR$ was).

| | U | | | | H | | | |
|---|---|---|---|---|---|---|---|---|
| | $\langle\Delta\rangle_L$ (unit) | $\langle\Delta\rangle_L$ (z) | $p$ | $n_L$ | $\langle\Delta\rangle_L$ (unit) | $\langle\Delta\rangle_L$ (z) | $p$ | $n_L$ |
| $\delta$ ($\mu V^2$) | 24.769 | 1.816 | 0.0005 | 4 | −8.725 | −1.181 | 0.0021 | 7 |
| $\theta$ ($\mu V^2$) | 5.030 | 1.869 | 0.0002 | 4 | −1.547 | −1.665 | 0.0003 | 7 |
| $\alpha$ ($\mu V^2$) | 11.176 | 2.329 | 0.0002 | 4 | −0.597 | −1.406 | 0.0004 | 7 |
| $\beta$ ($\mu V^2$) | 20.278 | 1.558 | 0.0005 | 4 | −1.812 | −1.731 | 0.0003 | 7 |
| $\gamma$ ($\mu V^2$) | 2.180 | 0.717 | 0.1404 | 4 | −1.947 | −2.421 | 0.0003 | 7 |
| $ap_{exp}$ | 0.217 | 2.440 | 0.0002 | 4 | 0.040 | 0.611 | 0.0930 | 7 |
| $ap_{off}$ ($\log \mu V^2$) | 0.350 | 2.032 | 0.0002 | 4 | −0.103 | −1.111 | 0.0021 | 7 |
| $\sum SMR$ ($\mu V^2$) | 27.956 | 1.832 | 0.0005 | 4 | 1.989 | 0.690 | 0.0601 | 7 |

**Table S3: Pretrial lesion sensitivity statistics**. Pretrial $\langle\Delta\rangle_L$ is shown original units and in $z$-score (as reported in the text) alongside $p$-values from permutation testing. $n_L$ refers to the number of lesions where this feature was observed before/after the lesion (oscillatory peaks were not always identified on each recording day, although aggregate SMR power $\sum SMR$ was).

| | U | | | H | | |
|---|---|---|---|---|---|---|
| | $\rho$ | $p$ | $N$ | $\rho$ | $p$ | $N$ |
| $\delta$ | −0.013 | 0.9209 | 384 | −0.026 | 0.5702 | 672 |
| $\theta$ | −0.039 | 0.9209 | 384 | −0.075 | 0.2063 | 672 |
| $\alpha$ | −0.015 | 0.9209 | 384 | 0.068 | 0.2063 | 672 |
| $\beta$ | −0.015 | 0.9209 | 384 | 0.026 | 0.5702 | 672 |
| $\gamma$ | −0.000 | 0.9975 | 384 | −0.050 | 0.3596 | 672 |
| $ap_{exp}$ | −0.050 | 0.9209 | 384 | 0.047 | 0.3596 | 672 |
| $ap_{off}$ | −0.049 | 0.9209 | 384 | −0.013 | 0.7392 | 672 |
| $\sum SMR$ | −0.014 | 0.9209 | 384 | 0.135 | 0.0038 | 672 |

**Table S4: Spatial correlation statistics**. Spearman correlation between $z$-scored first-difference induced by lesions on each single-channel spectral features ($\Delta$) and Manhattan distance to nearest lesion electrode.

| | U | | | H | | |
|---|---|---|---|---|---|---|
| | $SSM_{LL}$ | Null 95% PI | $p$ | $SSM_{LL}$ | Null 95% PI | $p$ |
| Behavior | $-1.13 \pm 0.00$ | [−1.18, −1.14] | 0.012 | $-0.87 \pm 0.00$ | [−0.88, −0.82] | 0.876 |
| Periodogram | $-0.24 \pm 0.00$ | [−0.32, −0.29] | 0.001 | $-0.28 \pm 0.00$ | [−0.37, −0.33] | 0.001 |
| Tensor | $-0.82 \pm 0.00$ | [−0.89, −0.85] | 0.001 | $-0.75 \pm 0.00$ | [−0.83, −0.76] | 0.014 |

**Table S5: Marginal log-likelihoods for LG-SSM models.** Mean $\pm$ SD (across initializations) of mean marginal log-likelihood for each dataset ($SSM_{LL}$) is shown alongside the 95 % percentile interval for the permuted null models.

| | | U | | | H | | |
|---|---|---|---|---|---|---|---|
| | | $d$ (unit) | $d$ (z) | $p$ | $d$ (unit) | $d$ (z) | $p$ |
| $T_{rxn}$ $(ms)$ | 1 | $14.64 \pm 0.80$ | $1.92 \pm 0.10$ | 0.030 | $-4.08 \pm 0.16$ | $-0.17 \pm 0.01$ | 0.824 |
| | 2 | $5.38 \pm 0.45$ | $0.70 \pm 0.06$ | 0.645 | $-1.32 \pm 0.15$ | $-0.05 \pm 0.01$ | 0.824 |
| | 3 | $-0.42 \pm 0.39$ | $-0.06 \pm 0.05$ | 0.956 | $-6.78 \pm 0.17$ | $-0.28 \pm 0.01$ | 0.824 |
| | 4 | $0.72 \pm 0.31$ | $0.09 \pm 0.04$ | 0.956 | $-0.96 \pm 0.31$ | $-0.04 \pm 0.01$ | 0.824 |
| | 5 | $0.20 \pm 0.19$ | $0.03 \pm 0.02$ | 0.956 | $-2.27 \pm 0.18$ | $-0.09 \pm 0.01$ | 0.824 |
| $v_{reach}$ $(cm/s)$ | 1 | $-0.40 \pm 0.07$ | $-0.25 \pm 0.04$ | 0.783 | $-0.58 \pm 0.11$ | $-0.14 \pm 0.03$ | 0.824 |
| | 2 | $-0.77 \pm 0.04$ | $-0.49 \pm 0.02$ | 0.726 | $-0.68 \pm 0.10$ | $-0.17 \pm 0.02$ | 0.824 |
| | 3 | $0.50 \pm 0.04$ | $0.32 \pm 0.03$ | 0.783 | $-0.83 \pm 0.11$ | $-0.21 \pm 0.03$ | 0.824 |
| | 4 | $0.16 \pm 0.05$ | $0.10 \pm 0.03$ | 0.956 | $-0.11 \pm 0.13$ | $-0.03 \pm 0.03$ | 0.824 |
| | 5 | $-0.48 \pm 0.05$ | $-0.31 \pm 0.03$ | 0.783 | $-0.11 \pm 0.12$ | $-0.03 \pm 0.03$ | 0.824 |
| $\theta_{err}$ (deg) | 1 | $0.30 \pm 0.08$ | $0.27 \pm 0.07$ | 0.783 | $-2.96 \pm 0.21$ | $-0.34 \pm 0.02$ | 0.824 |
| | 2 | $-0.83 \pm 0.10$ | $-0.75 \pm 0.09$ | 0.645 | $-5.18 \pm 0.19$ | $-0.59 \pm 0.02$ | 0.824 |
| | 3 | $-0.54 \pm 0.11$ | $-0.49 \pm 0.10$ | 0.783 | $-3.66 \pm 0.21$ | $-0.42 \pm 0.02$ | 0.824 |
| | 4 | $-1.07 \pm 0.12$ | $-0.96 \pm 0.11$ | 0.525 | $-4.03 \pm 0.25$ | $-0.46 \pm 0.03$ | 0.824 |
| | 5 | $-0.39 \pm 0.12$ | $-0.35 \pm 0.11$ | 0.783 | $-1.17 \pm 0.22$ | $-0.13 \pm 0.02$ | 0.824 |

**Table S6: LG-SSM input parameter $d$ for behavior dataset.** Mean $\pm$ SD (across initializations) for the input parameters $d_1, d_2, \ldots, d_5$ are shown in original and $z$-scored units for the behavior dataset, alongside computed $p$-values from permutation tests.

| | | U | | | H | | |
|---|---|---|---|---|---|---|---|
| | | $d$ (unit) | $d$ (z) | $p$ | $d$ (unit) | $d$ (z) | $p$ |
| $ap_{exp}$ | 1 | $0.05 \pm 0.00$ | $1.36 \pm 0.10$ | 0.020 | $0.06 \pm 0.01$ | $0.62 \pm 0.13$ | 0.174 |
| | 2 | $0.04 \pm 0.00$ | $1.10 \pm 0.08$ | 0.032 | $0.08 \pm 0.01$ | $0.85 \pm 0.13$ | 0.080 |
| | 3 | $0.04 \pm 0.00$ | $1.01 \pm 0.09$ | 0.080 | $0.06 \pm 0.01$ | $0.66 \pm 0.12$ | 0.174 |
| | 4 | $0.01 \pm 0.00$ | $0.30 \pm 0.12$ | 0.570 | $0.03 \pm 0.01$ | $0.30 \pm 0.09$ | 0.555 |
| | 5 | $0.02 \pm 0.01$ | $0.41 \pm 0.17$ | 0.460 | $-0.00 \pm 0.01$ | $-0.01 \pm 0.08$ | 0.948 |
| $ap_{off}$ ($\log \mu V^2$) | 1 | $0.01 \pm 0.01$ | $0.09 \pm 0.16$ | 0.888 | $-0.05 \pm 0.02$ | $-0.27 \pm 0.10$ | 0.370 |
| | 2 | $0.00 \pm 0.01$ | $0.07 \pm 0.11$ | 0.888 | $-0.01 \pm 0.02$ | $-0.04 \pm 0.11$ | 0.886 |
| | 3 | $0.02 \pm 0.01$ | $0.27 \pm 0.13$ | 0.650 | $-0.00 \pm 0.02$ | $-0.01 \pm 0.08$ | 0.990 |
| | 4 | $-0.01 \pm 0.01$ | $-0.13 \pm 0.19$ | 0.888 | $-0.03 \pm 0.01$ | $-0.14 \pm 0.07$ | 0.725 |
| | 5 | $0.03 \pm 0.01$ | $0.48 \pm 0.25$ | 0.460 | $-0.04 \pm 0.01$ | $-0.20 \pm 0.06$ | 0.555 |
| $\sum SMR$ ($\mu V^2$) | 1 | $5.49 \pm 0.72$ | $1.20 \pm 0.16$ | 0.020 | $2.19 \pm 0.23$ | $0.95 \pm 0.10$ | 0.011 |
| | 2 | $5.85 \pm 0.58$ | $1.28 \pm 0.13$ | 0.020 | $0.92 \pm 0.22$ | $0.40 \pm 0.10$ | 0.370 |
| | 3 | $3.08 \pm 0.34$ | $0.67 \pm 0.08$ | 0.304 | $1.13 \pm 0.16$ | $0.49 \pm 0.07$ | 0.286 |
| | 4 | $3.21 \pm 0.38$ | $0.70 \pm 0.08$ | 0.240 | $0.63 \pm 0.14$ | $0.27 \pm 0.06$ | 0.527 |
| | 5 | $0.99 \pm 0.50$ | $0.22 \pm 0.11$ | 0.687 | $0.17 \pm 0.15$ | $0.07 \pm 0.07$ | 0.886 |
| $\delta$ ($\mu V^2$) | 1 | $-3.55 \pm 1.59$ | $-0.40 \pm 0.18$ | 0.460 | $-12.72 \pm 1.39$ | $-0.74 \pm 0.08$ | 0.036 |
| | 2 | $-2.80 \pm 1.18$ | $-0.32 \pm 0.13$ | 0.563 | $-1.45 \pm 1.79$ | $-0.08 \pm 0.10$ | 0.888 |
| | 3 | $0.92 \pm 1.08$ | $0.10 \pm 0.12$ | 0.851 | $-2.68 \pm 1.08$ | $-0.16 \pm 0.06$ | 0.791 |
| | 4 | $-3.54 \pm 1.48$ | $-0.40 \pm 0.17$ | 0.483 | $-3.57 \pm 1.23$ | $-0.21 \pm 0.07$ | 0.586 |
| | 5 | $1.48 \pm 1.99$ | $0.17 \pm 0.23$ | 0.743 | $-4.37 \pm 1.27$ | $-0.25 \pm 0.07$ | 0.487 |
| $\theta$ ($\mu V^2$) | 1 | $-1.17 \pm 0.13$ | $-0.80 \pm 0.09$ | 0.080 | $-1.71 \pm 0.26$ | $-0.64 \pm 0.10$ | 0.087 |
| | 2 | $-1.10 \pm 0.11$ | $-0.75 \pm 0.07$ | 0.153 | $-0.88 \pm 0.31$ | $-0.33 \pm 0.11$ | 0.527 |
| | 3 | $-0.85 \pm 0.13$ | $-0.58 \pm 0.09$ | 0.388 | $-0.97 \pm 0.23$ | $-0.36 \pm 0.09$ | 0.527 |
| | 4 | $-0.45 \pm 0.17$ | $-0.31 \pm 0.12$ | 0.630 | $-0.71 \pm 0.22$ | $-0.26 \pm 0.08$ | 0.555 |
| | 5 | $0.72 \pm 0.23$ | $0.49 \pm 0.15$ | 0.388 | $-0.16 \pm 0.22$ | $-0.06 \pm 0.08$ | 0.886 |
| $\alpha$ ($\mu V^2$) | 1 | $2.19 \pm 0.22$ | $1.17 \pm 0.12$ | 0.050 | $-0.67 \pm 0.04$ | $-1.07 \pm 0.07$ | 0.008 |
| | 2 | $2.40 \pm 0.22$ | $1.28 \pm 0.12$ | 0.050 | $-0.57 \pm 0.05$ | $-0.92 \pm 0.08$ | 0.011 |
| | 3 | $1.19 \pm 0.23$ | $0.64 \pm 0.12$ | 0.388 | $-0.21 \pm 0.04$ | $-0.34 \pm 0.07$ | 0.527 |
| | 4 | $0.84 \pm 0.30$ | $0.45 \pm 0.16$ | 0.460 | $-0.13 \pm 0.04$ | $-0.20 \pm 0.07$ | 0.690 |
| | 5 | $0.88 \pm 0.38$ | $0.47 \pm 0.20$ | 0.460 | $-0.46 \pm 0.03$ | $-0.74 \pm 0.05$ | 0.087 |
| $\beta$ ($\mu V^2$) | 1 | $1.67 \pm 0.34$ | $0.57 \pm 0.12$ | 0.415 | $-1.12 \pm 0.10$ | $-0.99 \pm 0.09$ | 0.008 |
| | 2 | $2.91 \pm 0.36$ | $1.00 \pm 0.12$ | 0.154 | $-1.17 \pm 0.11$ | $-1.03 \pm 0.10$ | 0.008 |
| | 3 | $1.57 \pm 0.36$ | $0.54 \pm 0.12$ | 0.460 | $-0.64 \pm 0.07$ | $-0.56 \pm 0.06$ | 0.271 |
| | 4 | $1.87 \pm 0.46$ | $0.64 \pm 0.16$ | 0.388 | $-0.71 \pm 0.06$ | $-0.63 \pm 0.06$ | 0.143 |
| | 5 | $1.92 \pm 0.58$ | $0.66 \pm 0.20$ | 0.388 | $-0.55 \pm 0.06$ | $-0.49 \pm 0.06$ | 0.174 |
| $\gamma$ ($\mu V^2$) | 1 | $-1.66 \pm 0.10$ | $-1.89 \pm 0.11$ | 0.020 | $-1.37 \pm 0.09$ | $-1.28 \pm 0.09$ | 0.008 |
| | 2 | $-1.10 \pm 0.08$ | $-1.25 \pm 0.09$ | 0.040 | $-1.31 \pm 0.11$ | $-1.23 \pm 0.10$ | 0.008 |
| | 3 | $-1.03 \pm 0.08$ | $-1.17 \pm 0.09$ | 0.053 | $-0.91 \pm 0.05$ | $-0.85 \pm 0.05$ | 0.020 |
| | 4 | $-0.40 \pm 0.09$ | $-0.46 \pm 0.10$ | 0.460 | $-0.66 \pm 0.04$ | $-0.62 \pm 0.04$ | 0.129 |
| | 5 | $-0.07 \pm 0.11$ | $-0.08 \pm 0.12$ | 0.888 | $-0.31 \pm 0.06$ | $-0.29 \pm 0.06$ | 0.508 |

**Table S7: LG-SSM input parameter $d$ for periodogram dataset.** Mean $\pm$ SD (across initializations) for the input parameters $d_1, d_2, \ldots, d_5$ are shown in original and $z$-scored units for the periodogram dataset, alongside computed $p$-values from permutation tests.

| | | U | | | H | | |
|---|---|---|---|---|---|---|---|
| | | $d$ (unit) | $d$ (z) | $p$ | $d$ (unit) | $d$ (z) | $p$ |
| $\tau_1^d$ (a.u.) | 1 | $-0.05 \pm 0.00$ | $-1.78 \pm 0.10$ | 0.013 | $-0.03 \pm 0.00$ | $-0.85 \pm 0.05$ | 0.015 |
| | 2 | $-0.04 \pm 0.00$ | $-1.54 \pm 0.11$ | 0.013 | $-0.03 \pm 0.00$ | $-0.80 \pm 0.03$ | 0.052 |
| | 3 | $-0.01 \pm 0.00$ | $-0.47 \pm 0.09$ | 0.572 | $-0.02 \pm 0.00$ | $-0.42 \pm 0.05$ | 0.323 |
| | 4 | $-0.00 \pm 0.00$ | $-0.11 \pm 0.10$ | 0.872 | $-0.01 \pm 0.00$ | $-0.38 \pm 0.05$ | 0.323 |
| | 5 | $0.01 \pm 0.00$ | $0.35 \pm 0.13$ | 0.585 | $-0.01 \pm 0.00$ | $-0.39 \pm 0.07$ | 0.323 |
| $\tau_2^d$ (a.u.) | 1 | $0.05 \pm 0.00$ | $1.38 \pm 0.14$ | 0.013 | $-0.01 \pm 0.00$ | $-0.34 \pm 0.04$ | 0.323 |
| | 2 | $0.05 \pm 0.00$ | $1.38 \pm 0.12$ | 0.020 | $-0.01 \pm 0.00$ | $-0.32 \pm 0.05$ | 0.323 |
| | 3 | $0.02 \pm 0.00$ | $0.57 \pm 0.09$ | 0.420 | $-0.01 \pm 0.00$ | $-0.19 \pm 0.06$ | 0.493 |
| | 4 | $0.01 \pm 0.00$ | $0.27 \pm 0.12$ | 0.687 | $-0.00 \pm 0.00$ | $-0.08 \pm 0.06$ | 0.760 |
| | 5 | $0.00 \pm 0.01$ | $0.02 \pm 0.20$ | 0.968 | $-0.01 \pm 0.00$ | $-0.37 \pm 0.06$ | 0.323 |
| $\tau_3^d$ (a.u.) | 1 | $-0.01 \pm 0.00$ | $-0.67 \pm 0.11$ | 0.420 | $-0.02 \pm 0.00$ | $-0.62 \pm 0.02$ | 0.030 |
| | 2 | $0.00 \pm 0.00$ | $0.49 \pm 0.07$ | 0.572 | $-0.03 \pm 0.00$ | $-0.73 \pm 0.02$ | 0.015 |
| | 3 | $0.00 \pm 0.00$ | $0.27 \pm 0.09$ | 0.858 | $-0.01 \pm 0.00$ | $-0.38 \pm 0.03$ | 0.323 |
| | 4 | $0.00 \pm 0.00$ | $0.07 \pm 0.14$ | 0.941 | $-0.01 \pm 0.00$ | $-0.34 \pm 0.04$ | 0.306 |
| | 5 | $-0.00 \pm 0.00$ | $-0.02 \pm 0.24$ | 0.976 | $-0.00 \pm 0.00$ | $-0.11 \pm 0.04$ | 0.493 |
| $\tau_4^d$ (a.u.) | 1 | $0.06 \pm 0.01$ | $1.29 \pm 0.15$ | 0.040 | | | |
| | 2 | $0.04 \pm 0.01$ | $0.82 \pm 0.14$ | 0.300 | | | |
| | 3 | $0.00 \pm 0.01$ | $0.09 \pm 0.11$ | 0.968 | | | |
| | 4 | $0.01 \pm 0.01$ | $0.16 \pm 0.14$ | 0.872 | | | |
| | 5 | $-0.01 \pm 0.01$ | $-0.21 \pm 0.24$ | 0.872 | | | |

**Table S8: LG-SSM input parameter *d* for tensor dataset.** Mean $\pm$ SD (across initializations) for the input parameters $d_1, d_2, \ldots, d_5$ are shown in original and $z$-scored units for the tensor dataset, alongside computed $p$-values from permutation tests.

| | $T_{rxn}$ | | $v_{reach}$ | | $\theta_{err}$ | | $N$ |
|---|---|---|---|---|---|---|---|
| | $\rho$ | $p$ | $\rho$ | $p$ | $\rho$ | $p$ | |
| $\delta$ | 0.031 | 0.8182 | 0.027 | 0.8513 | 0.115 | 0.3690 | 100 |
| $\theta$ | −0.044 | 0.7723 | −0.146 | 0.1800 | 0.149 | 0.2987 | 100 |
| $\alpha$ | 0.409 | 0.0001 | −0.359 | 0.0011 | 0.348 | 0.0030 | 100 |
| $\beta$ | 0.394 | 0.0002 | −0.299 | 0.0071 | 0.339 | 0.0030 | 100 |
| $\gamma$ | 0.064 | 0.6681 | −0.245 | 0.0326 | −0.052 | 0.6548 | 100 |
| $ap_{exp}$ | 0.191 | 0.0993 | −0.193 | 0.0846 | 0.090 | 0.4354 | 100 |
| $ap_{off}$ | 0.111 | 0.3773 | −0.144 | 0.1800 | 0.113 | 0.3690 | 100 |
| $SMR_{\downarrow}^{P}$ | 0.326 | 0.0027 | −0.229 | 0.0381 | 0.284 | 0.0148 | 100 |
| $SMR_{\uparrow}^{P}$ | 0.177 | 0.1206 | −0.387 | 0.0005 | −0.145 | 0.2987 | 100 |
| $SMR_{\downarrow}^{f}$ | −0.416 | 0.0001 | 0.388 | 0.0005 | −0.184 | 0.1881 | 100 |
| $SMR_{\uparrow}^{f}$ | −0.426 | 0.0001 | 0.235 | 0.0370 | −0.336 | 0.0030 | 100 |
| $SMR_{\downarrow}^{W}$ | −0.238 | 0.0342 | 0.175 | 0.1134 | −0.094 | 0.4354 | 100 |
| $SMR_{\uparrow}^{W}$ | 0.000 | 1.0000 | 0.000 | 1.0000 | 0.000 | 1.0000 | 100 |
| $\sum SMR$ | 0.304 | 0.0050 | −0.327 | 0.0031 | 0.119 | 0.3690 | 100 |
| $\beta_{ref}$ | 0.350 | 0.0005 | −0.234 | 0.0448 | 0.322 | 0.0020 | 100 |
| $MRBD$ | −0.263 | 0.0097 | 0.000 | 0.9974 | −0.311 | 0.0023 | 100 |
| $PMBR$ | −0.441 | 0.0000 | 0.305 | 0.0090 | −0.365 | 0.0007 | 100 |
| $\tau_1^d$ | −0.378 | 0.0002 | 0.182 | 0.0971 | −0.175 | 0.0944 | 100 |
| $\tau_2^d$ | 0.418 | 0.0000 | −0.298 | 0.0090 | 0.321 | 0.0020 | 100 |
| $\tau_3^d$ | 0.150 | 0.1367 | −0.118 | 0.2821 | 0.101 | 0.3166 | 100 |
| $\tau_4^d$ | 0.507 | 0.0000 | −0.219 | 0.0507 | 0.410 | 0.0002 | 100 |

**Table S9: Behavioral correlation statistics for Monkey U.** Spearman correlation ($\rho$) alongside corresponding $p$-values were computed using differenced spectral/behavioral longitudinal time series. $N$ indicates the number of first differences that were used to compute each correlation.

| | $T_{rxn}$ | | $v_{reach}$ | | $\theta_{err}$ | | $N$ |
|---|---|---|---|---|---|---|---|
| | $\rho$ | $p$ | $\rho$ | $p$ | $\rho$ | $p$ | |
| $\delta$ | 0.072 | 0.6609 | 0.297 | 0.1064 | 0.320 | 0.0435 | 65 |
| $\theta$ | 0.145 | 0.4222 | 0.342 | 0.0741 | 0.415 | 0.0042 | 65 |
| $\alpha$ | 0.368 | 0.0355 | 0.262 | 0.1064 | 0.284 | 0.0514 | 65 |
| $\beta$ | 0.330 | 0.0510 | 0.241 | 0.1064 | 0.248 | 0.0812 | 65 |
| $\gamma$ | 0.024 | 0.8466 | 0.234 | 0.1064 | 0.298 | 0.0450 | 65 |
| $ap_{exp}$ | 0.112 | 0.5264 | 0.162 | 0.2504 | −0.003 | 0.9810 | 65 |
| $ap_{off}$ | 0.080 | 0.6609 | 0.245 | 0.1064 | 0.167 | 0.2577 | 65 |
| $SMR^{P}_{\downarrow}$ | 0.287 | 0.2253 | −0.066 | 0.7651 | −0.248 | 0.1382 | 48 |
| $SMR^{P}_{\uparrow}$ | 0.055 | 0.7181 | −0.240 | 0.1064 | −0.260 | 0.0790 | 63 |
| $SMR^{f}_{\downarrow}$ | −0.219 | 0.3141 | 0.229 | 0.1831 | 0.357 | 0.0450 | 48 |
| $SMR^{f}_{\uparrow}$ | 0.215 | 0.2535 | 0.176 | 0.2338 | 0.073 | 0.6675 | 63 |
| $SMR^{W}_{\downarrow}$ | 0.179 | 0.4222 | 0.004 | 0.9772 | 0.045 | 0.8202 | 48 |
| $SMR^{W}_{\uparrow}$ | 0.141 | 0.4222 | −0.007 | 0.9772 | −0.123 | 0.4308 | 63 |
| $\sum SMR$ | 0.217 | 0.2535 | −0.240 | 0.1064 | −0.425 | 0.0042 | 65 |
| $\beta_{ref}$ | −0.274 | 0.0549 | −0.155 | 0.3281 | 0.041 | 0.7484 | 65 |
| $MRBD$ | 0.226 | 0.1057 | 0.181 | 0.2994 | 0.066 | 0.7238 | 65 |
| $PMBR$ | 0.621 | 0.0000 | 0.312 | 0.0676 | 0.093 | 0.6883 | 65 |
| $\tau^{d}_{1}$ | −0.016 | 0.8970 | −0.011 | 0.9292 | 0.096 | 0.6883 | 65 |
| $\tau^{d}_{2}$ | 0.366 | 0.0081 | 0.279 | 0.0728 | 0.284 | 0.1326 | 65 |
| $\tau^{d}_{3}$ | 0.175 | 0.1956 | 0.085 | 0.5983 | 0.133 | 0.6883 | 65 |

**Table S10: Behavioral correlation statistics for Monkey H.** Spearman correlation ($\rho$) alongside corresponding $p$-values were computed using differenced spectral/behavioral longitudinal time series. $N$ indicates the number of first differences that were used to compute each correlation.

[3] IE Bray, SE Clarke, KM Casey, and P Nuyujukian. Neuroelectrophysiology-Compatible Electrolytic Lesioning. eLife. 2024;12:RP84385.

[4] A Tor, SE Clarke, IE Bray, P Nuyujukian, and tBI Laboratory. Material Damage to Multielectrode Arrays after Electrolytic Lesioning Is in the Noise. eLife. 2025;14.

[19] S Espenhahn, HE Rossiter, BCM van Wijk, N Redman, JM Rondina, J Diedrichsen, and NS Ward. Sensorimotor Cortex Beta Oscillations Reflect Motor Skill Learning Ability after Stroke. Brain Communications. 2020;2:fcaa161.

[37] CE Lunneborg. Random Assignment of Available Cases: Let the Inferences Fit the Design. 2000.

[38] EB Page. Ordered Hypotheses for Multiple Treatments: A Significance Test for Linear Ranks. Journal of the American Statistical Association. 1963;58:216–230.

[39] Y Benjamini and Y Hochberg. Controlling the False Discovery Rate: A Practical and Powerful Approach to Multiple Testing. Journal of the Royal Statistical Society. Series B (Methodological). 1995;57:289–300.

[40] S Seabold and J Perktold. Statsmodels: Econometric and Statistical Modeling with Python. scipy. 2010.
